# Endogenous serine proteases in MDCK sublines support trypsin-independent influenza A virus replication

**DOI:** 10.64898/2026.11.16.766931

**Authors:** Emma Balibrea Roelans, Jia-Yi Chan, Lisa Butticaz, Etori Aguiar Moreira, Gert Zimmer

## Abstract

The haemagglutinin (HA) of influenza A viruses (IAV) must be proteolytically cleaved by trypsin-like proteases to mediate fusion between the viral envelope and endosomal membrane of the host cell, a crucial step in virus entry. Most established cell lines rely on exogenous trypsin for the multicycle replication of influenza A viruses. In this study, we investigated various sublines of Madin-Darby canine kidney (MDCK) cells and found that MDCK-NBL2, MDCK-I, and MDCK-II support trypsin-independent replication of various influenza A viruses, including human, porcine, and avian strains. The serine protease inhibitors camostat and nafamostat mesylate reduced viral replication in a dose-dependent manner. However, the inhibitory effect diminished with increasing viral dose, most likely due to competition between the HA and the inhibitors for binding on the protease. We detected the endogenous expression of several transmembrane serine proteases in MDCK sublines, including TMPRSS2 and TMPRSS4. The characterization of these two canine serine proteases showed that TMPRSS4, but not TMPRSS2, is mainly localizing to the plasma membrane. Furthermore, canine TMPRSS4 was more effective than TMPRSS2 in terms of proteolytic activation of various influenza viruses, although there were strain-dependent differences. These results indicate that certain MDCK sublines express functional HA-activating proteases, providing useful models for the replication of influenza viruses without exogenous trypsin, as well as for the evaluation of antiviral strategies targeting transmembrane serine proteases.

**Importance:** The HA of most influenza viruses contains a monobasic cleavage site that is recognized and processed by cellular transmembrane serine proteases. MDCK cells are frequently used for the propagation and titration of influenza viruses, as well as for the production of influenza vaccines. We demonstrate that several MDCK sublines express various transmembrane serine proteases that support trypsin-independent replication of IAV. TMPRSS4 exhibits particularly high proteolytic activity at the cell surface and is accessible to membrane-impermeable serine protease inhibitors such as camostat and nafamostat mesylate. Furthermore, this protease is capable of activating various IAV strains. These properties suggest that it is necessary to investigate this protease, which has received little attention to date, in greater detail, including in other species, such as humans.

## Introduction

The haemagglutinin (HA) is the major envelope glycoprotein of influenza A viruses (IAV) and is essential for the initial stages of the infection. HA mediates attachment of the virion to host cells by binding to sialic acid-containing receptors on the plasma membrane and facilitates pH-dependent membrane fusion (1). After the virus is internalized via receptor-mediated endocytosis, acidification of the endosomal lumen by the vacuolar H^+^-ATPase (V-ATPase) induces a conformational change in HA, triggering fusion of the viral envelope with the endosomal membrane and release of the viral ribonucleoproteins into the cytosol (2). For HA to mediate fusion, the precursor protein HA_0_ must be proteolytically cleaved at a specific site to generate the HA_1_ and HA_2_ subunits (3). HA_1_ contains the receptor-binding domain, while the membrane-anchored HA_2_ subunit harbors the fusion machinery. The newly generated N-terminus of HA_2_ forms a highly conserved hydrophobic fusion peptide, which is initially buried within the HA structure but becomes exposed upon endosomal acidification, allowing it to insert into the target membrane to initiate membrane fusion (4).

The amino acid sequence at the HA proteolytic cleavage site of HA is a key determinant of viral pathogenicity as it dictates which host proteases can cleave and activate HA (5). Highly pathogenic avian influenza (HPAI) viruses of subtypes H5 and H7 typically have a cleavage site characterized by multiple basic amino acids matching the consensus motif R-X-(K/R)-R, which is recognized by prohormone convertases such as furin in the *trans*-Golgi network (5, 6). Since furin is ubiquitously expressed and active in the secretory pathway, HA with a polybasic cleavage site can be activated in many tissues, enabling systemic viral replication and contributing to the severe, often fatal disease seen in HPAI-infected poultry (3, 7). In contrast, most avian and mammalian IAV have a monobasic HA cleavage site which is recognized by trypsin-like proteases that are restricted to certain tissues. As a result, HA activation and productive replication of these viruses are generally limited to specific tissues (3, 5). Avian IAVs with a monobasic HA cleavage site are typically associated with a low-pathogenic phenotype and are referred to as low-pathogenic avian influenza (LPAI) viruses (8).

Several trypsin-like proteases including TMPRSS2, TMPRSS4 and human airway trypsin-like protease (HAT; TMPRSS11D) can activate IAVs with a monobasic HA cleavage site and can support multicycle replication *in vitro* (9–11). TMPRSS2 and HAT have also been shown to activate influenza B viruses. TMPRSS2, which is expressed in the human respiratory tract, is considered a key cellular protease for the activation of several human IAV strains (9, 12). However, many established cell lines lack sufficient endogenous protease activity to efficiently activate the HA of IAVs with a monobasic cleavage site. As a result, exogenous proteases such as trypsin are commonly added to the cell culture medium to enable multicycle replication of these viruses (13–15).

Madin-Darby canine kidney (MDCK) cells are widely used for influenza virus isolation, titration, and vaccine production (15, 16). Although it has been reported that MDCK cells lack HA-activating activity and require genetic modification to express TMPRSS2 or HAT for trypsin-independent IAV replication (17), there is evidence suggesting that MDCK cells are heterogeneous in their expression of proteases capable of supporting replication of IAVs with a monobasic HA cleavage site. For example, MDCK(+) cells can support multicycle replication of various IAV strains without exogenous trypsin, whereas IAV replication completely depends on trypsin in MDCK(-) cells (18). Notably, both sublines support influenza B virus replication in the absence of trypsin (18). Similarly, MDCK-NBL2 cells have been shown to support HA cleavage and multicycle replication of IAV without trypsin (19). However, single-cell cloning revealed that the MDCK-NBL2 cell line is heterogenous. Some clones required exogenous trypsin for IAV replication, while others did not. The specific cellular proteases responsible for trypsin-independent replication in these cells remain unidentified.

In this study, we further investigated the ability of certain MDCK sublines to support trypsin-independent replication of IAVs, suggesting the presence of endogenous activating proteases. We examined various cell lines for proteolytic activity capable of activating different HA subtypes, used specific inhibitors to determine whether these proteases are serine proteases, and identified transcripts of several serine proteases in MDCK cells. Additionally, we characterized the subcellular localization and HA-activating properties of the canine TMPRSS2 and TMPRSS4 proteases, providing new insights into the determinants of IAV cell tropism and the potential for trypsin-independent virus propagation in cell culture.

## Results

### MDCK cells support trypsin-independent replication of H1N1_PR8_

The trypsin-independent replication of A/Puerto Rico/8/1934 (H1N1_PR8_) was analysed in several established cell lines, including various Madin-Darby canine kidney (MDCK) cell sublines. MDCK-NBL2 cells were isolated from the normal kidney tissue of an adult female Cocker Spaniel and deposited in September 1958 by S.H. Madin and N.B. Darby at the American Type Culture Collection (ATCC, CCL-34). We also included the MDCK-I and MDCK-II sublines, which differ in several characteristics, including transepithelial electrical resistance (20), glycosphingolipid composition (21), expression of the mucin-type membrane protein gp40 (22), and susceptibility to bat influenza viruses (23). The MDCK-MR subline was included as it has been reported to lack trypsin-independent influenza virus replication (17). Human colon carcinoma Caco-2 cells, known to support trypsin-independent IAV propagation via endogenous TMPRSS2 and TMPRSS4 expression (24), served as a positive control. We also included human lung adenocarcinoma cell lines Calu-3 and A549, and human lung fibroblasts (MRC-5) in our analysis.

The cell lines were inoculated with H1N1_PR8_ at a multiplicity of infection (MOI) of 0.0001 and incubated for 72 h in fetal bovine serum (FBS)-deficient cell culture medium in absence or presence of 1.0 µg/mL of acetylated trypsin. At 24, 48, and 72 h post infection (p.i.), cell culture supernatants were collected, and infectious virus titers were determined by titration on MDCK-II cells (**Fig. 1**). In Caco-2 cells, H1N1_PR8_ replicated efficiently without trypsin, showing similar kinetics to those observed with trypsin (**Fig. 1A**). In MDCK-I cells, replication of H1N1_PR8_ without trypsin resulted in only slightly lower viral titers compared to those with exogenous trypsin (**Fig. 1B**), whereas in MDCK-II (**Fig. 1C**) and MDCK-NBL2 (**Fig. 1D**) cells, titers were substantially lower in the absence of trypsin. The MDCK-MR subline did not support trypsin-independent H1N1_PR8_ replication (**Fig. 1E**) and only minimal replication was observed in Calu-3 cells (**Fig. 1F**), despite reported TMPRSS2 and other serine protease expression (25, 26). No trypsin-independent replication of H1N1_PR8_ was detected in A549 (**Fig. 1G**) and MRC-5 (**Fig. 1H**) cells. These results indicate that the ability of cell lines to support trypsin-independent H1N1_PR8_ replication varies, likely reflecting differences in endogenous protease expression.

**Fig. 1.**
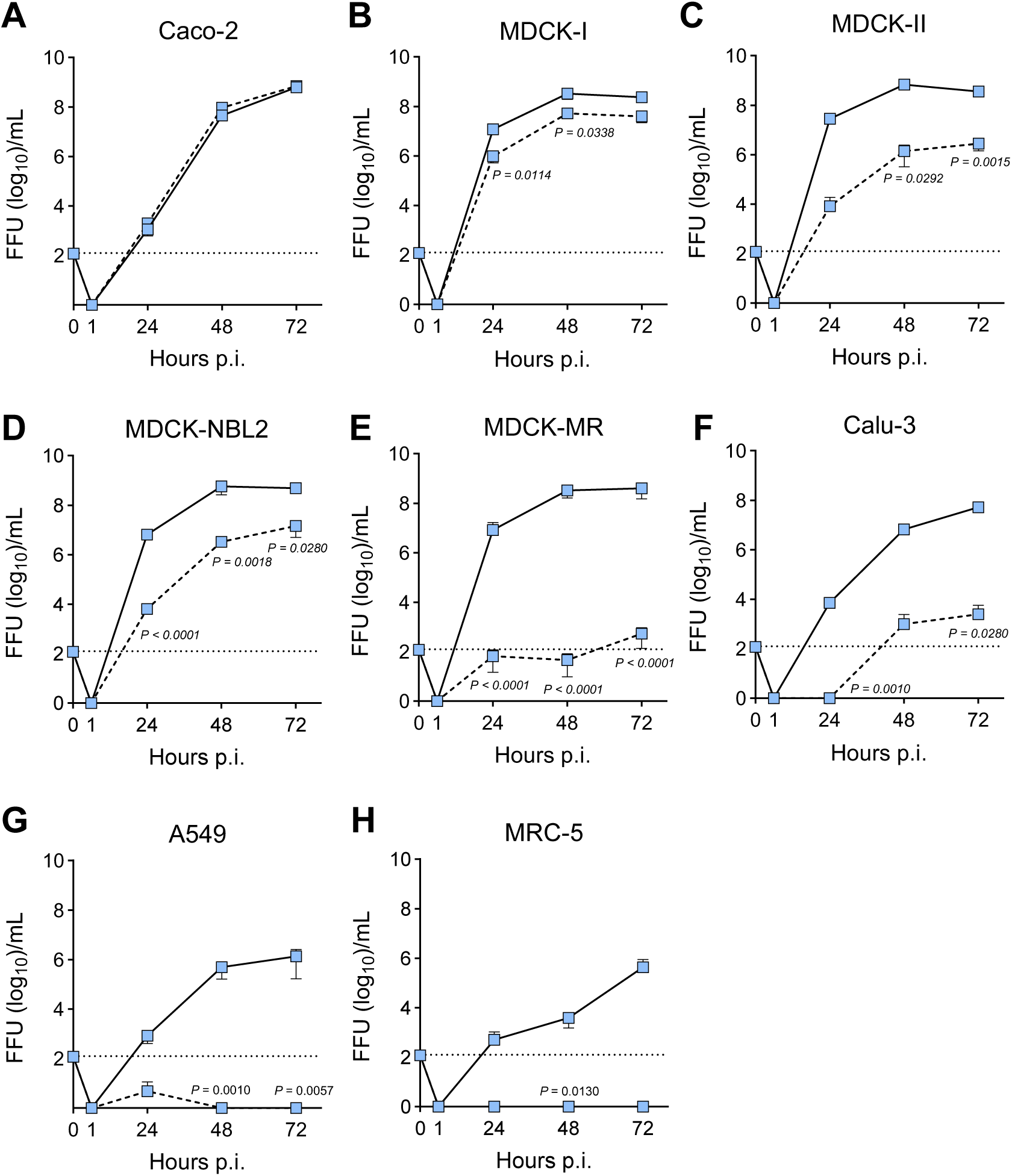
Trypsin-independent replication of H1N1_PR8_ on established cell lines. Caco-2 (**A**), MDCK-I (**B**), MDCK-II (**C**), MDCK-NBL2 (**D**), MDCK-MR (**E**), Calu-3 (**F**), A549 (**G**), and MRC-5 (**H)** cells were seeded into 6-well cell culture plates at a density of 250’000 cells/mL and infected with H1N1_PR8_ at an MOI of 0.0001. The cells were incubated for 72 h at 37 °C in FBS-free MEM medium either in the presence (continuous lines) or absence of acetylated trypsin (dashed lines). At the indicated time points, infectious virus titers in the cell culture supernatants were determined by endpoint dilution assay on MDCK-II cells. Data represent mean values ± SD from three biological replicates. Statistical significance was determined with two-way ANOVA with Geisser-Greenhouse’s correction and Šídák’s multiple comparisons test. Only statistically significant *P* values are shown in the graphs.

### MDCK-NBL2 cells support trypsin-independent replication of various IAV subtypes

To assess whether endogenous proteases in MDCK cells can activate IAVs strains beyond H1N1_PR8_, we analyzed a panel of viruses representing different HA subtypes and cleavage-site motifs (**Fig. 2**). H1N1_HH4/09_, which like H1N1_PR8_ contains a single arginine at the HA cleavage site, replicated in the absence of exogenous trypsin, although titers were consistently lower than with trypsin (**Fig. 3A**). The H3N2 viruses A/Udorn/307/1972 (H3N2_Ud/72_) and A/swine/Bakum/IDT1769/2003 (H3N2_Bak/03_), both with a **K**/**R**-Q-T-**R** motif (**Fig. 2**), showed delayed replication without trypsin (**Fig. 3B, C**). The LPAI virus A/duck/Potsdam/1402/1986 (H5N2_Po/86_), with an **R**-E-T-**R** motif, replicated efficiently without trypsin (**Fig. 3D**). In contrast, A/swan/Potsdam62/1981 (H7N7_Po/81_) and A/turkey/Wisconsin/1/1966 (H9N2_Wis/66_), with P-**K**-G-**R** and V-S-S-**R** motifs, respectively, exhibited inefficient trypsin-independent replication (**Fig. 3E, F**).

**Fig. 2.**
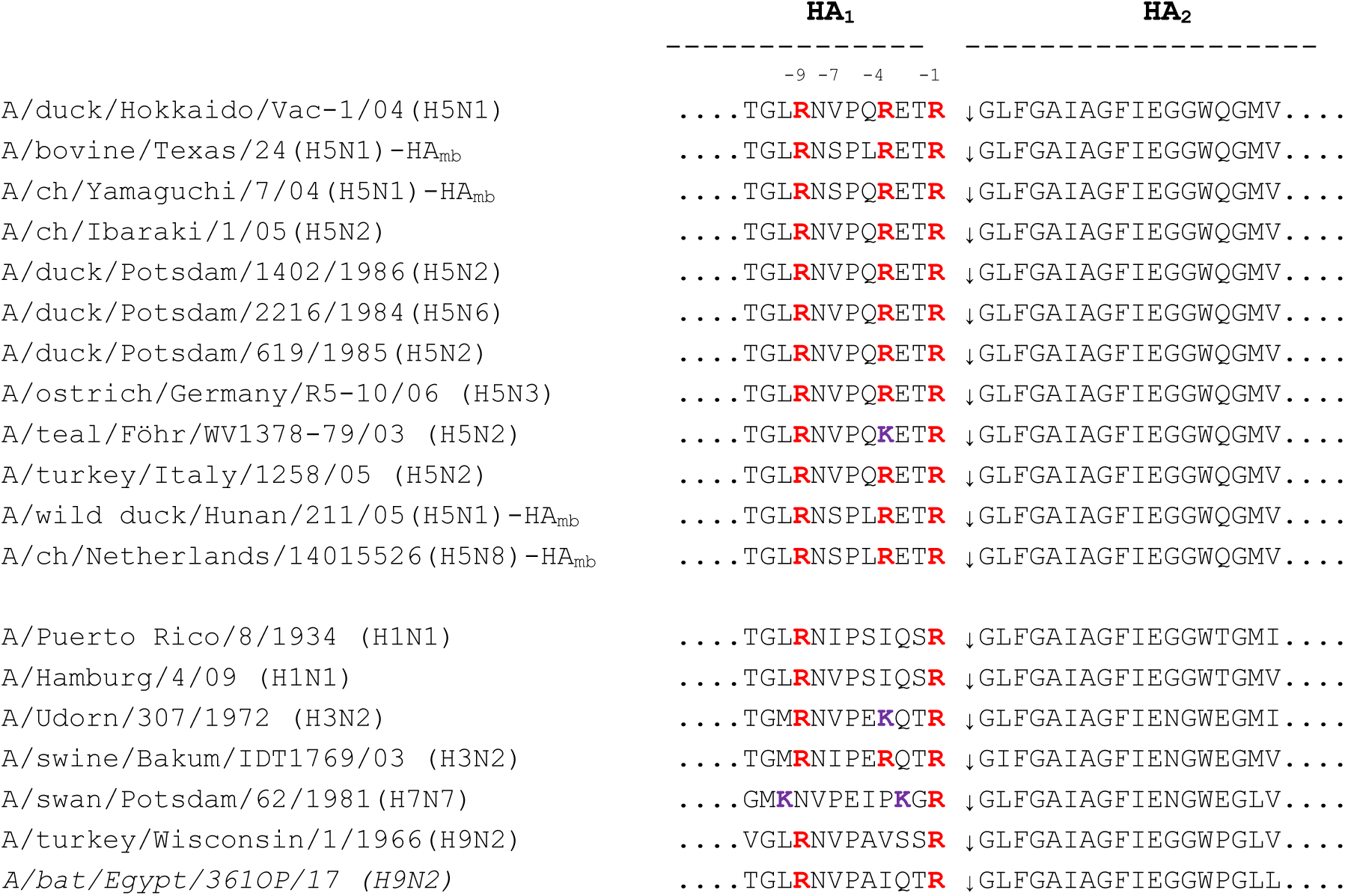
Amino acid sequences at the proteolytic cleavage site in the HA of various IAVs. Amino acid sequences are shown at which proteolytic cleavage into the HA_1_ and HA_2_ subunits occurs. The upper block depicts the cleavage sites for all H5Nx viruses used in this study, the sequences show the cleavage sites of viruses belonging to other subunits. The cleavage site is marked by an arrow. Arginine residues at the cleavage site and in its vicinity are shown in red letters, while lysine residues are shown in purple letters.

**Fig. 3.**
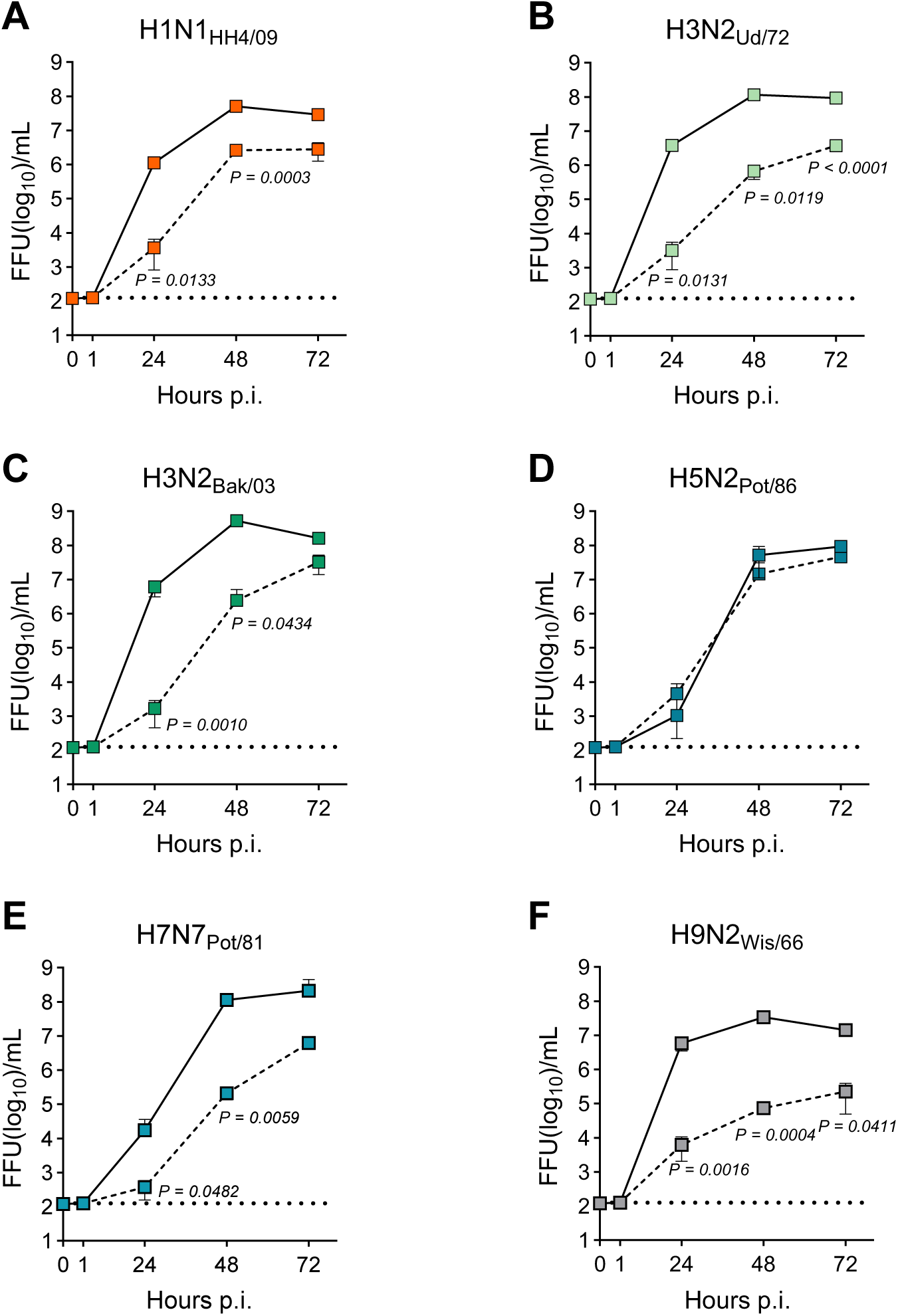
Trypsin-independent replication of IAV belonging to different subtypes in MDCK-NBL2 cells. MDCK-NBL2 cells that were grown in 6-well plates (approximately 10_6_ cells/well) were infected with H1N1_HH4/09_ (**A**), H3N2_Ud/72_ (**B**), H3N2_Bak/03_ (**C**), H5N2_Pot/86_ (**D**), H7N7_Pot/81_ (**E**), and H9N2_Wis/66_ (**F**) at an MOI of 0.0001. The cells were incubated for 72 h at 37 °C in FBS-free MEM medium either in the presence (continuous lines) or absence of acetylated trypsin (dashed lines). At the indicated time points, infectious virus titers in the cell culture supernatants were determined by endpoint dilution assay on MDCK-II cells. Data represent mean values ± SD from three biological replicates. Statistics were done with two-way ANOVA with Geisser-Greenhouse’s correction and Šídák’s multiple comparisons test.

To assess the efficiency of HA cleavage by endogenous serine proteases, we infected MDCK-NBL2 cells with H1N1_HH4_ at an MOI of 0.0001 and incubated them for 72 h with or without exogenous trypsin. Virus particles released into the cell culture supernatant were pelleted by ultracentrifugation, and viral proteins were analyzed by SDS-PAGE and immunoblotting with a polyclonal antibody against the 2009 pandemic H1N1 HA protein (**Fig. 4**). Under non-reducing conditions (-DTT), bands corresponding to monomeric, dimeric and trimeric HA were detected, but the precursor protein HA_0_ could not be distinguished from cleaved HA_1,2_ due to the intact disulfide bond. Under reducing conditions (+DTT), HA_0_, HA_1_, and HA_2_ were detected both in the presence and in the absence of trypsin. However, in the absence of trypsin, a substantial fraction of HA remained uncleaved, whereas HA_0_ was almost completely processed with trypsin. This reduced HA cleavage likely accounts for the lower infectious virus titers observed without trypsin (**Fig. 3A**). Overall, these results indicate that MDCK-NBL2 cells express endogenous proteases capable of activating IAV strains with multiple HA subtypes, though the efficiency of activation varies among strains.

**Fig. 4.**
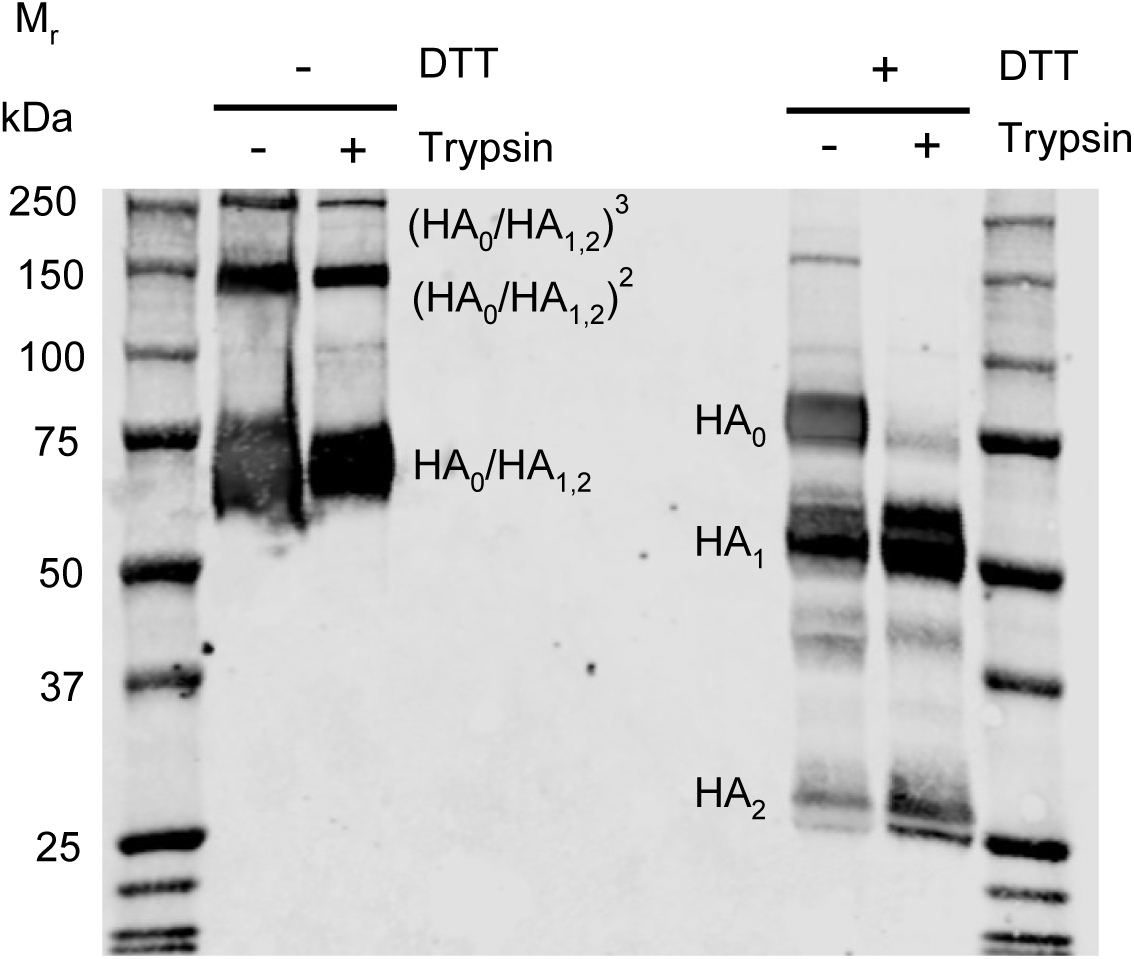
Western blot analysis of H1N1_HH4/09_ virions released from infected MDCK-NBL2 cells. MDCK-NBL2 cells were cultured in T75 cell culture flasks (approximately 10^7^ cells/flask) and infected with H1N1_HH4/09_ virus at a multiplicity of infection (MOI) of 0.0001. Following infection, cells were incubated at 37 °C in FBS-free MEM in the absence (−) or presence (+) of trypsin. At 72 h postinfection, cell culture supernatants were collected, and cell debris was removed by low-speed centrifugation. Viral particles were subsequently pelleted by ultracentrifugation. Virus pellets were resuspended in SDS sample buffer and subjected to SDS-PAGE under nonreducing (−DTT) or reducing (+DTT) conditions, followed by transfer to nitrocellulose membranes by semidry blotting. The HA protein of H1N1_HH4/09_ was detected with a polyclonal antibody against the HA protein of the 2009 pandemic H1N1 virus. Molecular masses (in kDa), corresponding to the migration of molecular mass markers, are indicated on the left. Bands corresponding to HA_0_, HA_1/2_, HA_1_, and HA_2_ are indicated. A representative result from two independent experiments is shown.

### MDCK-NBL2 cells support trypsin-independent replication of various H5Nx viruses

To investigate the extent to which different viral strains of the same HA subtype can replicate without exogenous trypsin, MDCK-NBL2 cells were infected with various LPAI H5Nx viruses, or with HPAI H5Nx viruses whose polybasic cleavage site had been modified to correspond to the monobasic cleavage site **R**-E-T-**R** (**Fig. 2**). Although all H5Nx strains possessed the same cleavage site motif, their ability to replicate efficiently in MDCK-NBL2 cells without trypsin varied considerably. The LPAI viruses A/duck/Potsdam/2216/1984 (H5N6_Po/84_) (**Fig. 5A**), A/duck/Potsdam/619/1985 (H5N2_Po/85_) (**Fig. 5B**) and A/ostrich/Germany/R5-10/2006 (H5N3_Ge/06_) (**Fig. 5C**) replicated nearly as well as without trypsin as with it. In contrast, LPAI strains A/duck/Hokkaido/Vac-1/2004 (H5N1_Hok/04_) (**Fig. 5D**), A/turkey/Italy/1258/2005 (H5N2_It/05_) (**Fig. 5E**) and A/teal/Föhr/WV1378-79/2003 (H5N2_Fö/03_) (**Fig. 5F**) showed significantly lower titers without trypsin. Similarly, the HPAI A/cattle/Texas/2024 (H5N1_Tex/24_), in which the polybasic cleavage site had been converted to a monobasic sequence motif, replicated much better with trypsin (**Fig. 5G**), while A/chicken/Yamaguchi/7/2004 (H5N1_Yam/04_) with the same motif replicated almost as efficiently without trypsin, though titers were lower at 24 h post infection (**Fig. 5H**).

**Fig. 5.**
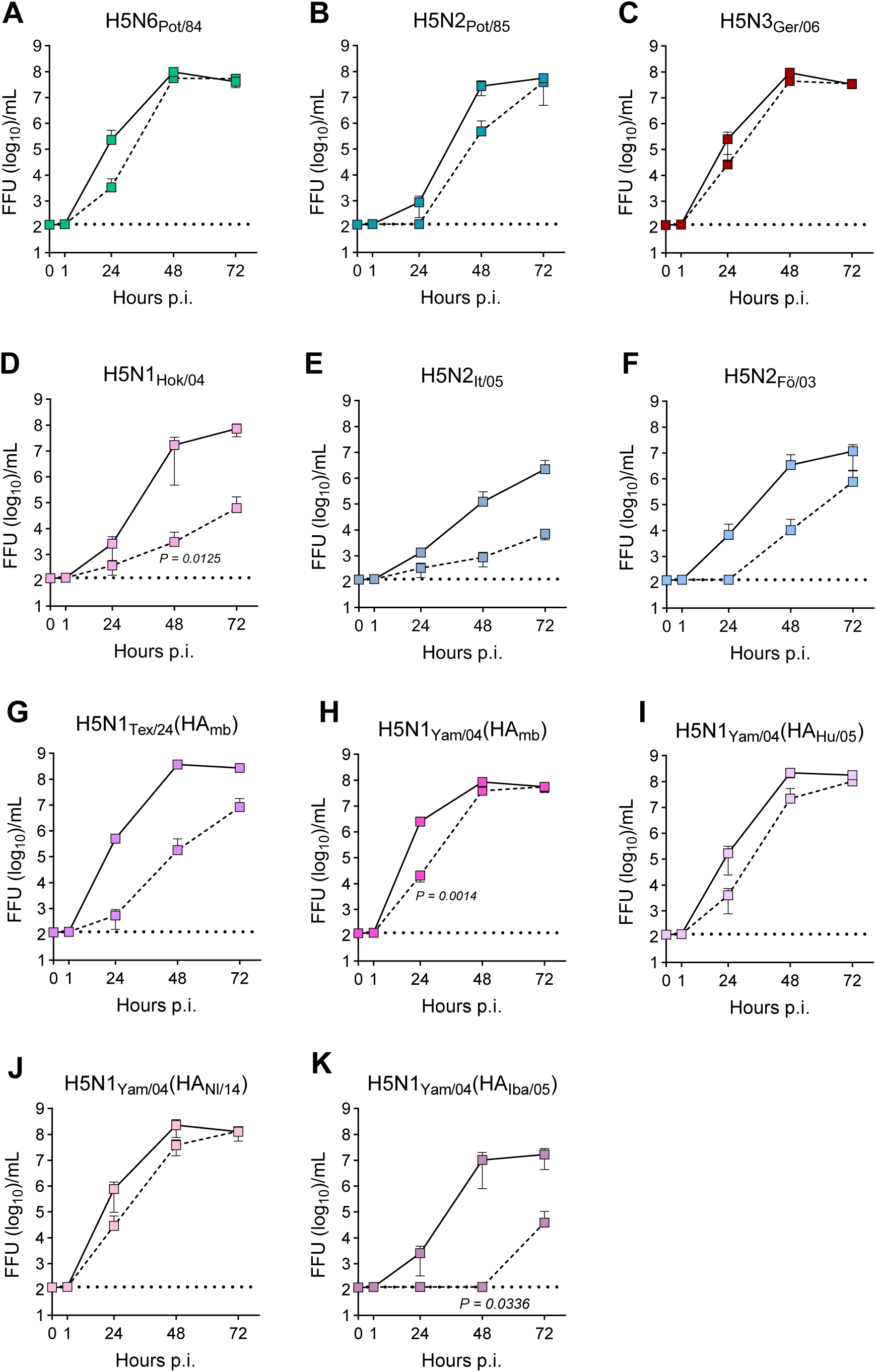
MDCK-NBL2 cells support trypsin-independent replication of various H5Nx viruses. MDCK-NBL2 cells grown in 6-well plates (approx. 10^6^ cells/well) were infected with H5N6_Pot/84_ (**A**), H5N2_Pot/85_ (**B**), H5N3_Ger/06_ (**C**), H5N1_Hok/04_ (**D**), H5N2_It/05_ (**E**), H5N2_Fö/03_ (**F**), H5N1_Tex/24_ (HA_mb_) (**G**), H5N1_Yam/04_ (HA_mb_) (**H**), H5N1_Yam/04_ (HA_Hu/05_) (**I**), H5N1_Yam/04_ (HA_Nl/14_) (**J**), and H5N1_Yam/04_ (HA_Iba/05_) (**K**) at an MOI of 0.0001 and incubated for 72 h at 37 °C in FBS-free MEM medium either in the presence (continuous lines) or absence of acetylated trypsin (dashed lines). At the indicated time points, infectious virus titers in the cell culture supernatants were determined by endpoint dilution assay on MDCK-II cells. Data represent mean values ± SD from three biological replicates. Statistics were done with two-way ANOVA with Šídák’s multiple comparisons test.

Efficient trypsin-independent replication was also observed for reassortant viruses based on H5N1_Yam/04_ with modified HA from A/wild duck/Hunan/211/2005 (**Fig. 5I**) or A/chicken/Netherlands/14015526 (**Fig. 5J**), whereas a reassortant virus with the HA from A/chicken/Ibaraki/1/2005 showed little replication without trypsin (**Fig. 5K**). These findings indicate that the amino acid sequence at the HA cleavage site alone does not determine efficient activation by endogenous proteases in MDCK-NBL2 cells.

### Serine protease inhibitors block trypsin-independent IAV replication in MDCK cells

Camostat mesylate, nafamostat mesylate, and AEBSF (4-(2-aminoethyl)benzenesulfonyl fluoride hydrochloride; Pefabloc) are synthetic serine protease inhibitors which inhibit a broad spectrum of serine proteases, although their inhibitory effect may vary depending on the protease (27). These inhibitors are water soluble and are expected to preferentially target serine proteases that are localized at the plasma membrane or are released into the extracellular environment. To determine whether the endogenous proteases expressed by MDCK cells are susceptible to these inhibitors, MDCK-NBL2 cells were infected with H1N1_PR8_ using an MOI of 0.0001 and incubated for 72 h in cell culture medium containing different concentrations of the inhibitors in the absence of exogenous trypsin. Cell culture supernatants were collected at 24, 48, and 72 h p.i. and infectious virus titers were determined.

Nafamostat mesylate inhibited H1N1_PR8_ replication in a concentration-dependent manner (**Fig. 6A**). Virus replication was completely inhibited at concentrations of 1 µM and 10 µM, whereas 0.1 µM had no significant effect. A dose-response analysis performed at 48 h p.i. showed that nafamostat mesylate reduced infectious virus titers by approximately 4 log_10_ over the concentration range of 0.5 to 1.0 µM (**Fig. 6B**). Camostat mesylate also inhibited H1N1_PR8_ replication in a concentration-dependent manner, completely preventing virus replication at concentrations of 10, 25, and 50 µM. At 1 µM, virus titers were significantly reduced but residual replication was still detectable (**Fig. 6C**). Dose-response analysis demonstrated an approximately 4 log_10_ reduction in infectious virus titers at camostat mesylate concentrations ranging from 1 to 5 µM (**Fig. 6D**), indicating that camostat mesylate was less potent than nafamostat mesylate against the endogenous serine protease activity in MDCK-NBL2 cells. AEBSF, an irreversible, broad-spectrum serine protease inhibitor (28), completely inhibited H1N1_PR8_ replication only at 100 µM, whereas concentrations of 1 and 10 µM had little or no inhibitory effect (**Fig. 6E**). Thus, among the three inhibitors tested, AEBSF was the least effective at inhibiting the endogenous protease activity required for H1N1_PR8_ replication in MDCK-NBL2 cells.

**Fig. 6.**
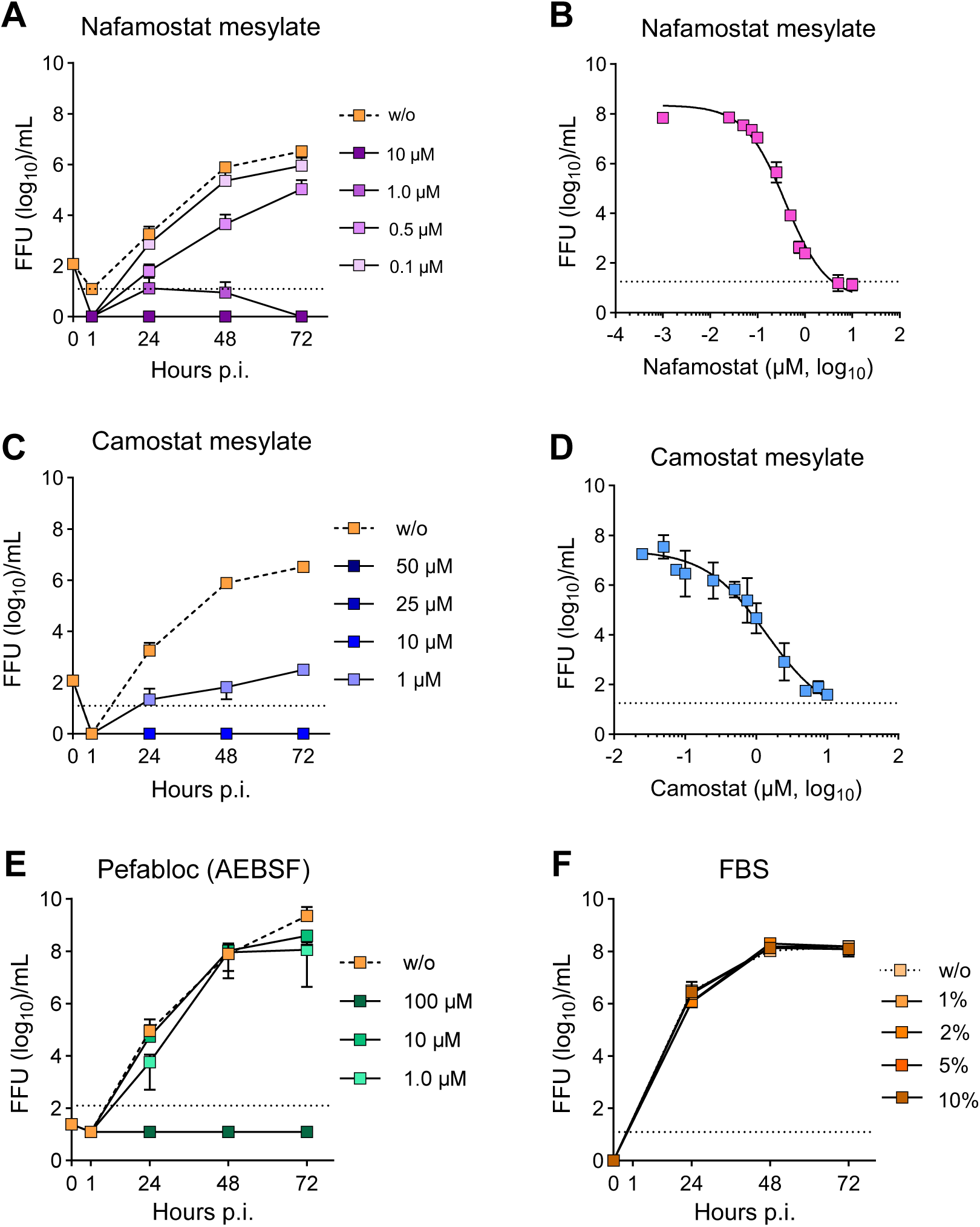
Serine protease inhibitors block trypsin-independent IAV replication in MDCK cells. (**A - F**) MDCK-NBL2 cells were inoculated with H1N1_PR8_ at an MOI of 0.0001. The cells were subsequently incubated at 37 °C for 72 h with FBS-free MEM containing the indicated concentrations of nafamostat mesylate (**A, B**), camostat mesylate (**C, D**), Pefabloc (**E**) or FBS (**F**). Aliquots of cell culture supernatant were collected at 24, 48 and 72 h post infection and subsequently titrated (**A**). Inhibitor dose-response curves were determined based on the virus titers obtained at 48 h post infection (**B, D**). Data represent mean values ± SD from three biological replicates. Statistics were done with two-way ANOVA with Geisser-Greenhouse’s correction and Šídák’s multiple comparisons test.

To investigate the effect of serine protease inhibitors as a function of viral dose, MDCK-NBL2 cells were infected with H1N1_PR8_, using MOIs ranging from 0.0001 to 1.0. The infected cells were incubated at 37 °C in the presence of 10 µM of either camostat mesylate or nafamostat mesylate, or in the absence of an inhibitor. The infectious titers in the cell culture supernatant were determined at 48 h post infection. When an MOI of 1.0 was used, the inhibitory effect by camostat mesylate was reduced by 36.6-fold while at lower virus doses the inhibitory effect was significantly higher, for example exceeding 500000 -fold when an MOI of 0.001 was used (**Fig. 7A, B**). A similar phenomenon was also observed for infected cells treated with 10 µM of nafamostat mesylate (**Fig. 7C, D**). These results provide further evidence that MDCK cells possess endogenous serine proteases that mediate trypsin-independent IAV replication. Serine protease inhibitors camostat and nafamostat efficiently prevent virus propagation on MDCK cells but, surprisingly, act in a virus dose-dependent manner although they are targeting a cellular and not a viral protein.

**Fig. 7.**
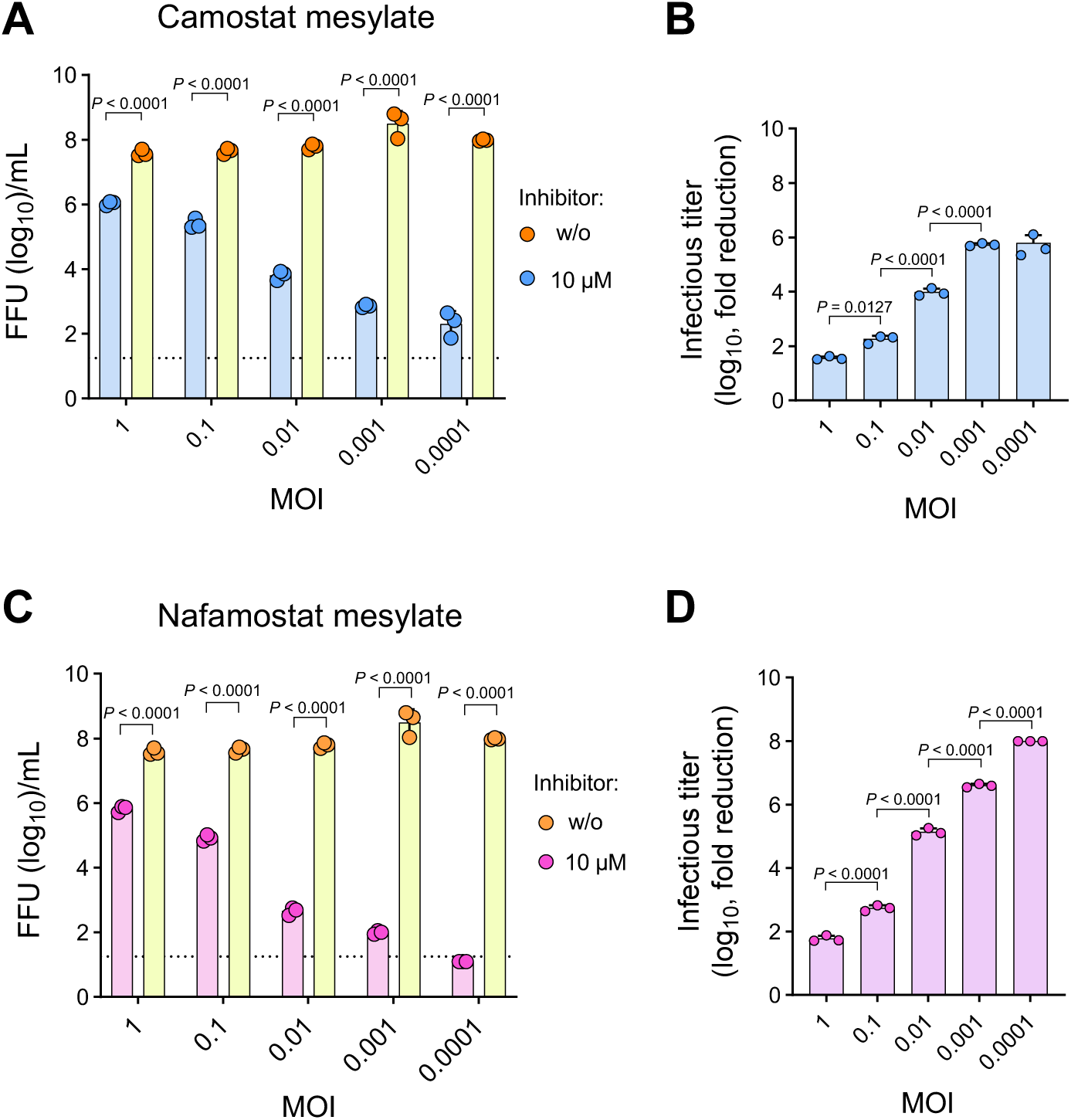
Effect of serine protease inhibitors on trypsin-independent H1N1_PR8_ replication in MDCK-NBL2 cells at different MOIs. **(A - D)** MDCK-NBL2 were infected with H1N1_PR8_ using the indicated MOIs. The infeceted cells were incubated for 48 h at 37 °C in FBS-deficient MEM containing 10 µM of camostat mesylate (**A, B**) or nafamostat mesylate (**C, D**). Aliquots of cell culture supernatant were collected at 48 h post infection (pi) and virus titers were determined. (**B, D**) The virus titer fold reduction was calculated for camostat mesylate (**B**) or nafamostat nemesylate (**D**). Data represent mean values ± SD from three biological replicates. Statistics were done with two-way ANOVA with Geisser-Greenhouse’s correction and Šídák’s multiple comparisons test.

### Identification of canine transmembrane serine proteases in MDCK cells

To date, there is no evidence that TMPRSS2 or other transmembrane serine proteases capable of activating influenza viruses are expressed by MDCK cells. To identify transcripts of canine TMPRSS family members, total RNA was isolated from MDCK-II and MDCK-NBL2 cells, and RT-PCR was performed using oligonucleotide primers designed to amplify approximately 500-bp cDNA fragments of canine TMPRSS genes (**Table 1**). Transcripts corresponding to TMPRSS1 (hepsin), TMPRSS2, TMPRSS4, TMPRSS6, TMPRSS11D (HAT, 650 bp), TMPRSS12, TMPRSS13, and TMPRSS14 (ST14, matriptase) were detected in both MDCK-NBL2 (**Fig. 8A**) and MDCK-II (**Fig. 8B**) cells. TMPRSS3 transcripts were detected in MDCK-II but not in MDCK-NBL2 cells. Transcripts corresponding to TMPRSS7, TMPRSS9, TMPRSS10, TMPRSS11A, and TMPRSS15 were not detected in either cell line. Although an approximately 300-bp product was observed in the RT-PCR for TMPRSS9, its identity is uncertain, because the expected amplicon size is 550-bp.

**Fig. 8.**
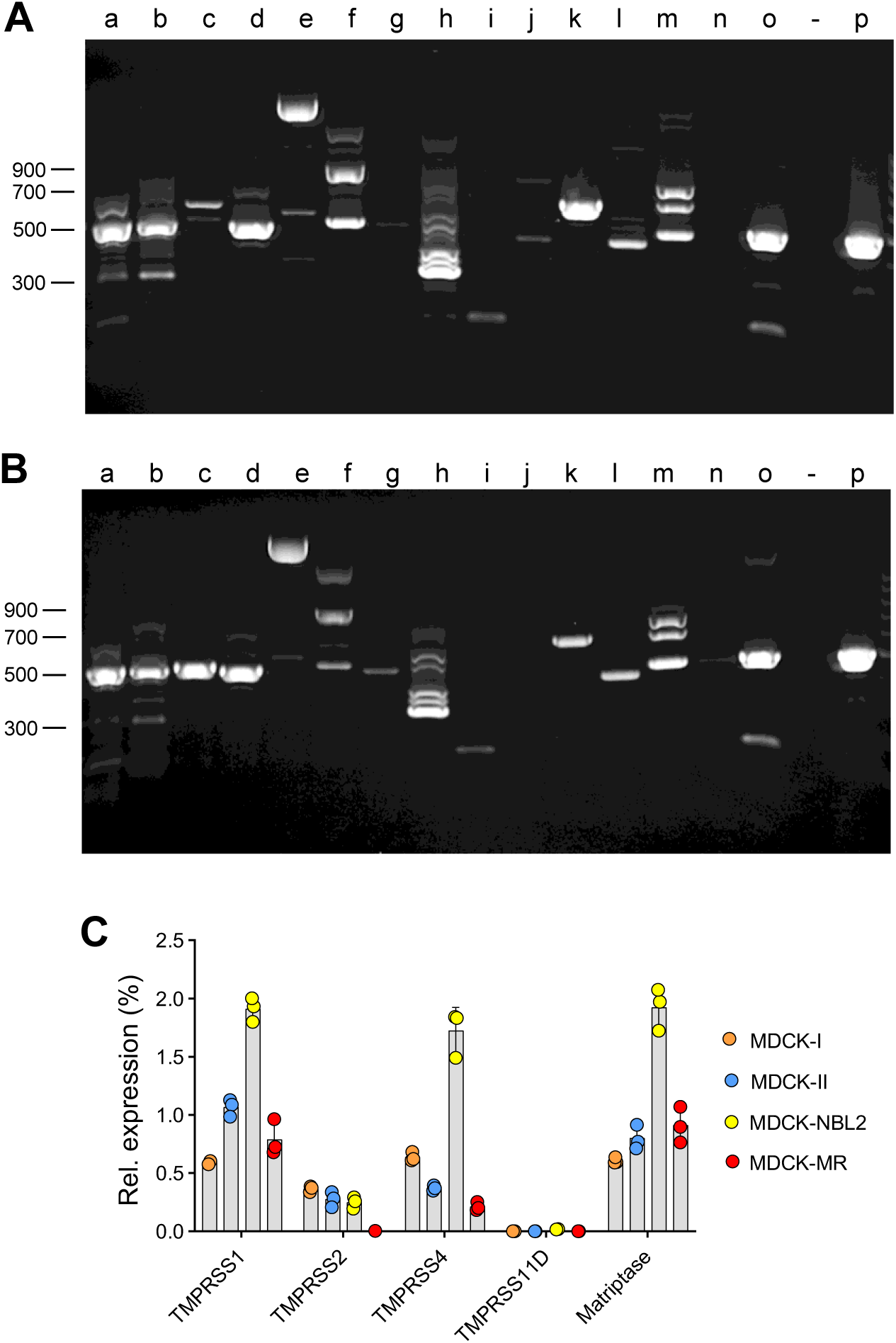
Identification of TMPRSS transcripts in MDCK cells. (**A, B**) Total RNA was extracted from MDCK-NBL2 (**A, C**) and MDCK-II (**B**) cells and reverse transcribed. Using specific oligonucleotide primers, cDNA fragments of approximately 500 base pairs (bp) were amplified for various canine TMPRSS proteases and GAPDH: TMPRSS1/hepsin (a), TMPRSS2 (b), TMPRSS3 (c), TMPRSS4 (d), TMPRSS5 (e), TMPRSS6 (f), TMPRSS7 (g), TMPRSS9 (h), TMPRSS10 (i), TMPRSS11A (j), TMPRSS11D/HAT (k), TMPRSS12 (l), TMPRSS13 (m), TMPRSS15 (n), ST14 (o), and GAPDH (p). On the left-hand side are molecular weight standards indicated. (**C**) Relative expression levels of TMPRSS proteases in four different MDCK cell lines. RT-qPCR was performed for the indicated TMPRSS proteases and the expression relative to the housekeeping gene GAPDH was calculated.

**Table 1.**
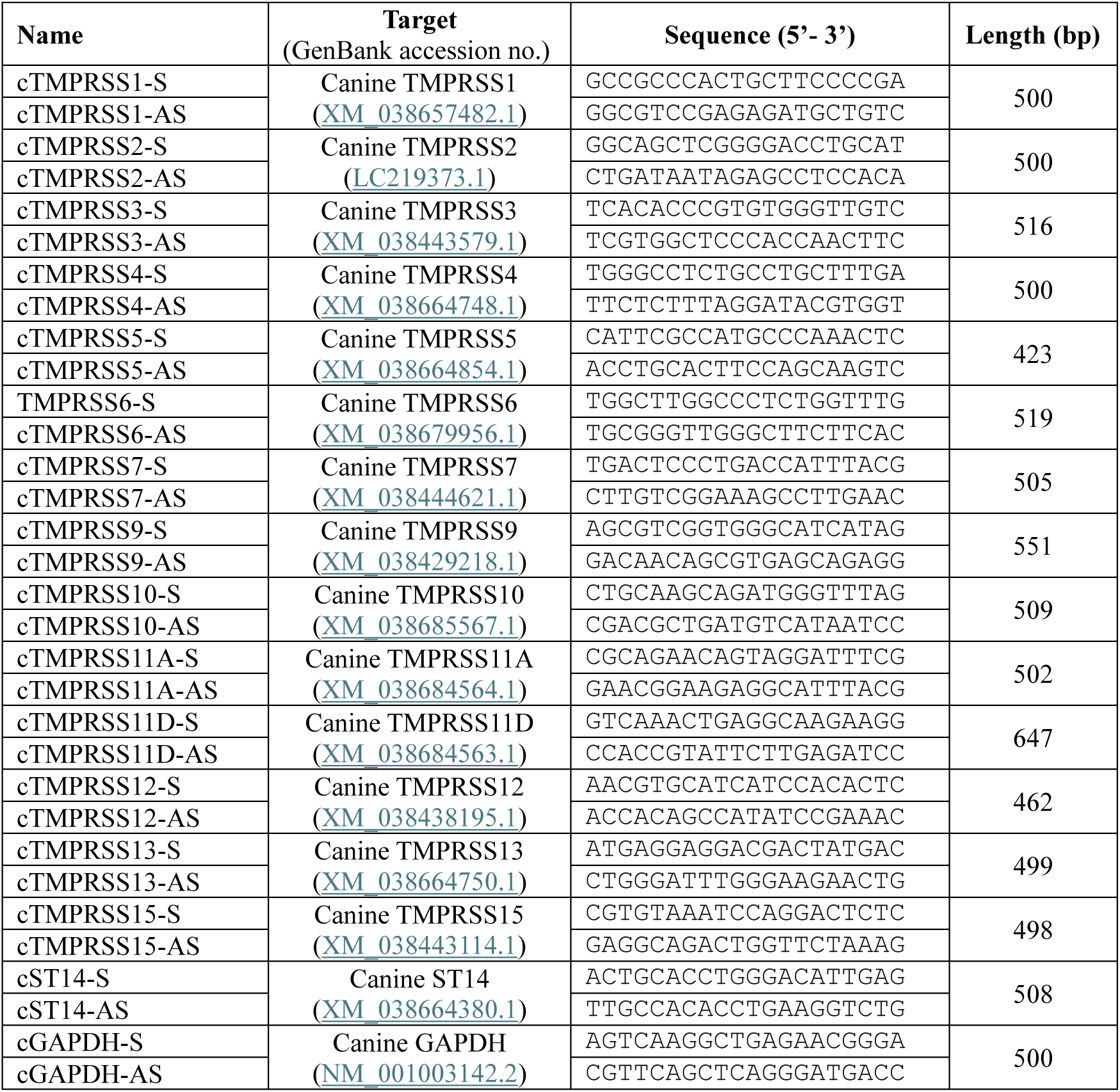
Oligonucleotide primers for the detection of TMPRSS transcripts by RT-PCR.

The cDNAs encoding the full-length open reading frames of the canine proteins TMPRSS1 (hepsin), TMPRSS2, TMPRSS4 and TMPRSS11D were obtained by RT-PCR using total RNA from MDCK-NBL2 cells and their sequences determined. A comparison of the predicted primary amino acid sequences showed that the proteins are identical or nearly identical to the published amino acid sequences derived from *Canis lupus familiaris* or *Canis lupus dingo* (**Table 2**). A comparison with the published sequences of the human orthologues of these transmembrane serine proteases revealed that the MDCK-TMPRSS1 protein is 94 per cent identical to its human orthologue while the other three proteases exhibit significantly lower levels of identity. Taking together, these results suggest that both various transmembrane serine proteases as well as cellular serine protease inhibitors are expressed by MDCK-NBL2 cells.

**Table 2.** Homology of transmembrane serine proteases and serine protease inhibitors of MDCK-NBL2 cells with their human counterparts.

| **MDCK-NBL2** |  | ***Canis lupus* orthologues** |  | **Human orthologues** |  |
|---|---|---|---|---|---|
| Protein | GenBank acc. | AA identity | GenBank acc. | AA identity | GenBank acc. |
| TMPRSS1 | QB150831 | 100% | XP_02577480 | 94% | NM_182983 |
| TMPRSS2 | QB150832 | 99.8% | BBD3386 | 80% | NM_001135099 |
| TMPRSS4 | QB150833 | 99.7% | XP_005619775 | 83% | NM_019894 |
| TMPRSS11D | QB150834 | 100% | XP_038540491 | 79% | NM_004262 |

To investigate whether MDCK cell lines differ in terms of TMPRSS transcription levels, the RNA transcripts of the proteases known to be capable of activating IAV (11, 12), were quantified in the four MDCK sublines using quantitative RT-qPCR, with the housekeeping gene GAPDH serving as reference (**Fig. 8C**). The MDCK sublines MDCK-NBL2, MDCK-I and MDCK-II exhibited significant levels of TMPRSS1, TMPRSS4 and matriptrase transcripts, but relatively low levels of TMPRSS2 (MDCK-I: 0.365%; MDCK-II: 0.276%; MDCK-NBL2: 0.249%) and very low levels of TMPRSS11D (MDCK-I: 0.004%; MDCK-II: 0.001%; MDCK-NBL2: 0.028%). MDCK-MR cells showed relatively high levels of TMPRSS1 (0.789%) and matriptase transcripts (0.911%), but only small amounts of TMPRSS4 (0.212%). Interestingly, no active transcription of TMPRSS2 or TMPRSS11D was detected in MDCK-MR cells.

### TMPRSS4 is expressed in its enzymatically active form at the plasma membrane

The endogenous concentration of transmembrane serine proteases in established cell lines is usually quite low, which makes their detection by antibodies difficult (29, 30). Furthermore, no antibodies were available against canine transmembrane proteases, some of which exhibit only moderate homology with their human orthologues (see **Table 2**). In order to investigate the subcellular localization of these proteins nonetheless, we cloned the cDNAs encoding the serine proteases cTMPRSS1, cTMPRSS2, cTMPRSS4, and cTMPRSS11D, derived from MDCK-NBL2 cells, into the pCMV-3Tag3 expression plasmid, such that the recombinant proteins were tagged with a triple FLAG epitope at their C-terminus. MDCK-MR cells were transfected with these plasmids, and the serine proteases detected by indirect immunofluorescence using a monoclonal antibody directed against the FLAG epitope. While all four transmembrane serine proteases could be detected using the FLAG antibody in intracellular compartments of fixed and detergent-permeabilized MDCK-MR cells, only cTMPRSS4 was also detectable on the surface of non-permeabilized cells (**Fig. 9A**). This finding suggests that cTMPRSS4 is transported to the plasma membrane more efficiently than the other proteases investigated, or that it stays at the plasma membrane for a longer time.

**Fig. 9.**
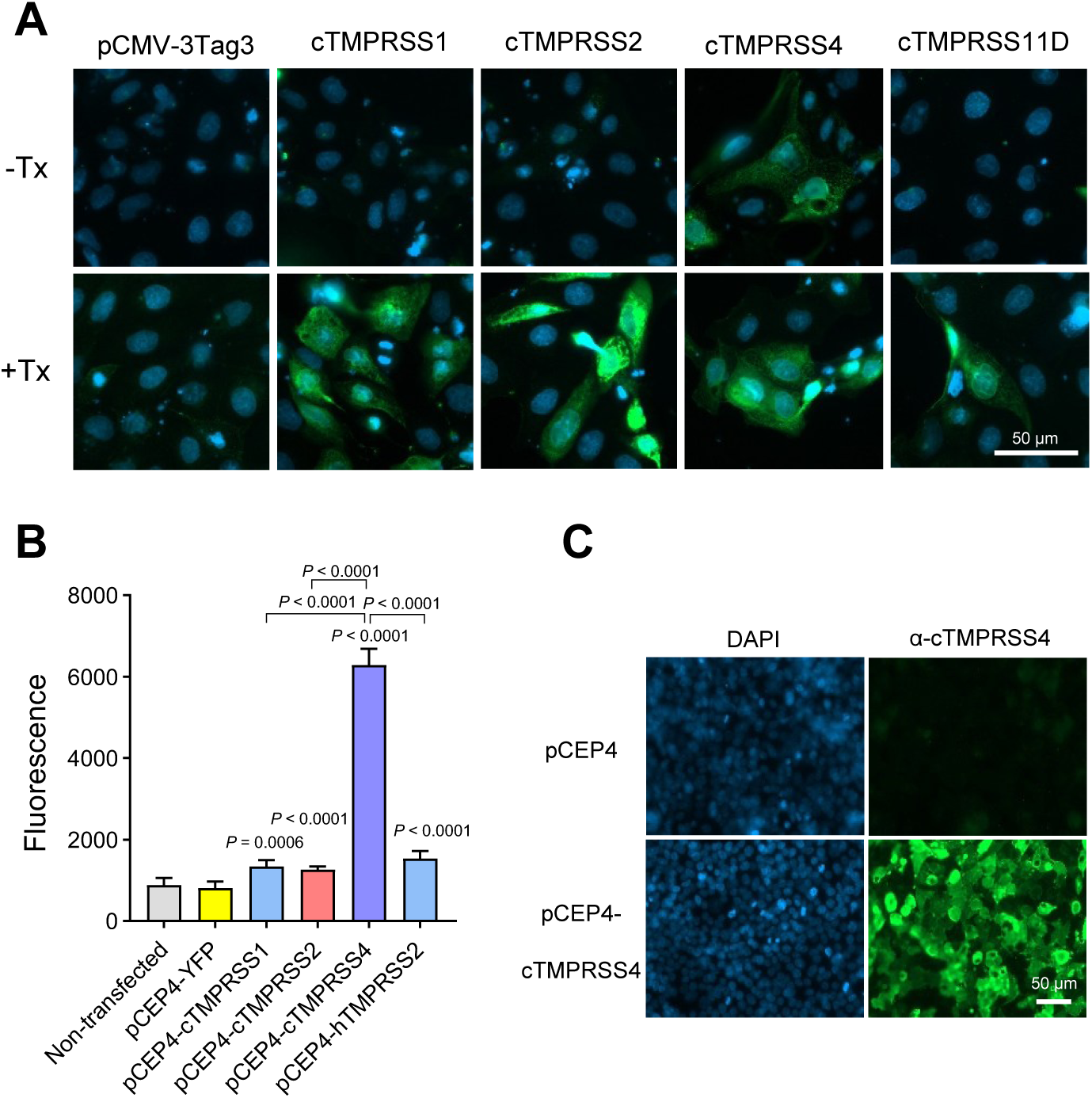
Canine TMPRSS4 is expressed in its enzymatically active form on the plasma membrane. (**A**) Indirect immunofluorescence analysis of MDCK-MR cells transfected with pCMV-3Tag3 plasmids encoding the specified canine transmembrane serine proteases. The cells were fixed with formalin 20 h after transfection and subsequently permeabilized with Triton X-100 (+Tx) or left non-permeabilized (-Tx). The serine proteases were detected using a monoclonal antibody directed against the triple FLAG epitope at the C terminus of the proteins (green fluorescence). The nuclei of the cells were stained with DAPI (blue fluorescence). The scale bar corresponds to 50 µm. (**B**) Assay of serine protease activity using a fluorogenic peptide substrate. BHK-21 cells were transfected with pCEP4 plasmids encoding the specified transmembrane serine proteases. Twenty hours after transfection, the cells were incubated with the Boc-QAR-AMC substrate and the fluorescence released after 2 hours at 37 °C was measured. The data represent mean values ± SD from eight biological replicates. Statistical analysis was performed using one-way ANOVA with Dunnett’s and Tukey’s tests for multiple comparisons. (**C**) Indirect immunofluorescence analysis of HEK-293T cells stably transfected with either an empty pCEP4 plasmid or pCEP4-cTMPRSS4. The formalin-fixed cells were stained with a chicken polyclonal anti-cTMPRSS4 serum chicken (green fluorescence) and nuclei were labelled with DAPI (blue fluorescence). The scale bar corresponds to 50 µm.

To assess the proteolytic activity of the transmembrane proteases on the cell surface, BHK-21 cells were transfected with expression plasmids encoding the canine serine proteases cTMPRSS1, cTMPRSS2, cTMPRSS4 or the human hTMPRSS2, without a FLAG epitope, to avoid any potential interference of this foreign epitope with enzyme activity. BHK-21 cells were chosen because they can be transfected with high efficiency and exhibit low endogenous serine protease activity. When the transfected cells were incubated for 2 h at 37 °C with the fluorogenic serine protease substrate Boc-QAR-AMC, a significant increase of the fluorescence signal was recorded for all 4 proteases compared to non-transfected cells or cells transfected with empty plasmid (**Fig. 9B**). However, the fluorescence signal released from cTMPRSS4-transfected cells was significantly higher than that produced by the other proteases. As the fluorogenic peptide is likely cell-impermeable, this finding suggests that cTMPRSS4 is expressed at the plasma membrane at higher levels than the other proteases investigated. Indirect immunofluorescence of stably transfected HEK-293T cells using a polyclonal immune serum raised against canine TMPRSS4 confirmed that this serine protease is expressed at the cell surface (**Fig. 9C**).

### Canine TMPRSS2 and TMPRSS4 activate influenza viruses in a virus strain-dependent manner

Human TMPRSS2 and TMPRSS4 have been shown to be key serine proteases facilitating trypsin-independent replication of H1N1_PR8_ in Caco-2 cells (24). To investigate whether the canine orthologues of these transmembrane serine proteases would show IAV-activating activity, we cloned the cDNAs encoding canine TMPRSS2 and TMPRSS4 into the pCEP4 vector plasmid which contains the EBNA-1 gene allowing episomal replication of the plasmid in human cells (31). HEK-293T cells were transfected with either empty pCEP4 plasmid, pCEP4-TMPRSS2 or pCEP4-TMPRSS4 and were subsequently selected with hygromycin B. We succeeded in generating stably transfected HEK-293T cells expressing either canine TMPRSS2, canine TMPRSS4, or no canine serine protease (pCEP4 control cells). To investigate whether the HEK-293T cells expressing either canine TMPRSS2 or TMPRSS4 would support trypsin-independent IAV replication, the cells were infected with various IAV at an MOI of 0.01 and incubated at 37 °C either in the presence or absence of exogenous trypsin. Infectious virus released into the cell culture supernatant was determined at 48 h post infection. HEK-293T cells stably transfected with empty pCEP4 plasmid supported IAV replication only if exogenous trypsin was added to the cell culture medium (**Fig. 10A-H**), in line with previous reports that these cells do not express endogenous proteases at levels sufficient to activate IAV (24). In HEK-293T cells stably transfected with canine TMPRSS2, both H1N1_PR8_ (**Fig. 10A**) and H1N1_HH4/09_ (**Fig. 10B**) replicated without trypsin to titers that were significantly lower than those in the presence of trypsin. In cells expressing canine TMPRSS4, both viruses showed a more efficient trypsin-independent replication than in HEK-293T-cTMPRSS2 cells. Likewise, H3N2_Bak/03_ replicated efficiently without trypsin in HEK-293T-cTMPRSS4 cells with only 7-fold lower titers compared to the condition with trypsin, whereas it was significantly less efficiently activated in HEK-293T-cTMPRSS2 cells (**Fig. 10C**). Interestingly, H3N2_Ud/72_ replicated in both HEK-293T-cTMPRSS2 and HEK-293T-cTMPRSS4 cells without trypsin to similar titers, however, these were 1,500-fold and 4,200-fold lower, respectively, compared to the condition with trypsin (**Fig. 10D**). This observation suggests that H3N2_Ud/72_ is less efficiently activated than H3N2_Bak/03_ in HEK-293T-cTMPRSS4 cells. HEK-293T-cTMPRSS2 cells supported H5N1_Yam/04_ (HA_mb_) replication without trypsin more efficiently than HEK-293T-cTMPRSS4 cells (**Fig. 10E**), while H5N1_Hok/04_ was inefficiently activated by either cell line (**Fig. 10F**). The avian H7N7_Po/81_ strain replicated inefficiently without trypsin in HEK-293T-cTMPRSS2 cells but efficiently in HEK-293T-cTMPRSS4 cells (**Fig. 10G**). HEK-293T-cTMPRSS2 cells did not at all support trypsin-independent replication of H9N2_Wis/66_, while HEK-293T-cTMPRSS4 cells allowed some replication of this virus strain without trypsin (**Fig. 10H**). These results suggest that the two canine transmembrane serine proteases differ significantly in terms of their ability to activate certain viral strains. However, each of the two proteases also exhibited preferences for specific viral strains, which were independent of the respective HA subtype.

**Fig. 10.**
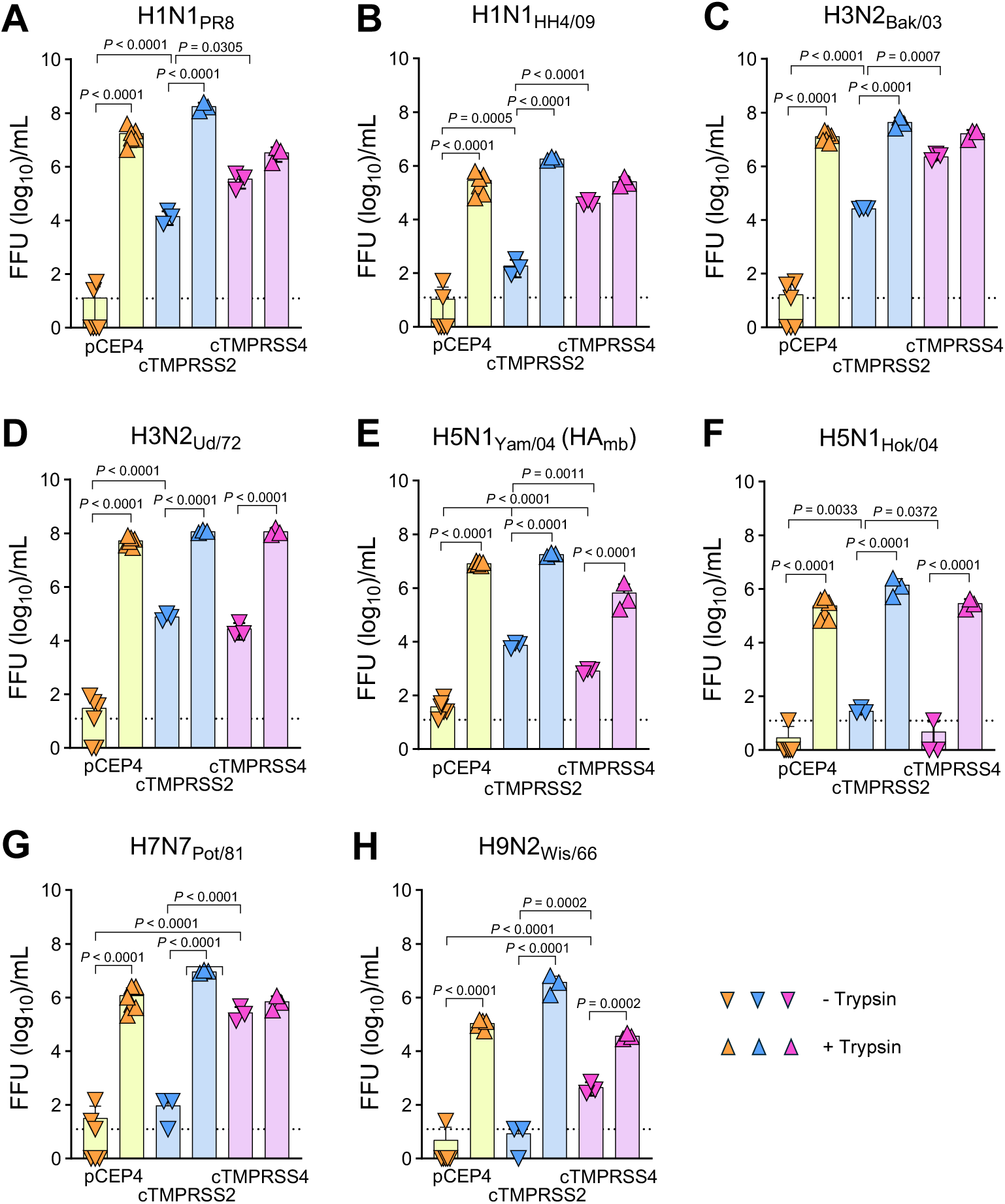
IAV activation by canine TMPRSS2 and TMPRSS4. HEK-293T cells stably transfected with either empty pCEP4 plasmid, recombinant pCEP4-cTMPRSS2 plasmid or recombinant pCEP4-cTMPRSS4 plasmid were infected with either H1N1_PR8_ (**A**), H1N1_HH4/09_ (**B**), H3N2_Bak/03_ (**C**), H3N2_Ud/72_ (**D**), H5N1_Yam/04_ (HA_mb_) (**E**), H5N1_Hok/04_ (**F**), H7N7_Pot/81_ (**G**) or H9N2_Wis/66_ (**H**) at an MOI of 0.01. The infected cells were incubated for 48 h in FBS-free medium with or without 0.4 µg/mL of acetylated trypsin. Infectious virus that was released into the cell culture supernatants at 48 h post infection was titrated on MDCK-II cells. Mean values and SD of three infection experiments are shown. Significant differences were assessed by the one-way ANOVA with Sidak’s multiple comparisons test.

## Discussion

It has previously been reported that MDCK cells lack IAV-activating proteases (17). When these cells were genetically modified to express human TMPRSS2 or HAT, IAV replication in the absence of trypsin was enabled (17). However, it has also been reported that certain MDCK sublines or clones are capable of supporting trypsin-independent IAV replication, suggesting that MDCK cells are heterogeneous regarding the expression of HA-activating proteases (18, 19). To date, it is not known which cellular proteases are responsible for this trypsin-independent replication in MDCK cells or how their expression profile differs among cell sublines or clones.

In this study, we demonstrate that MDCK-NBL2, MDCK-I, and MDCK-II but not MDCK-MR cells support trypsin-independent H1N1_PR8_ replication, which was effectively inhibited by the serine protease inhibitors camostat and nafamostat mesylate, indicating that HA-activating serine proteases are present in these MDCK cell sublines. Consistent with this observation, we detected the RNA transcripts of several transmembrane proteases in MDCK-I, MDCK-II, and MDCK-NBL2 cells, including TMPRSS1 (hepsin), TMPRSS2, TMPRSS4, TMPRSS11D (HAT), and TMPRSS14 (matriptase), the murine and human orthologs of which are known to proteolytically activate IAV (9, 10, 12, 24, 32–36).

Molecular cloning of cDNAs from MDCK-NBL2 cells, which encode the complete open reading frames of canine TMPRSS1, TMPRSS2, TMPRSS4 and TMPRSS11D, showed that the sequences are identical to the published canine sequences but exhibit only moderate homology with their human orthologues (see **Table 2**). Transient expression of the four transmembrane serine proteases, each tagged with a C-terminal FLAG epitope, showed that the proteins are predominantly localised in intracellular compartments; only canine TMPRSS4 was also detectable on the cell surface using the FLAG-specific antibody. Transient expression of the native proteases, which were not tagged with a FLAG epitope, followed by incubation of the transfected cells with a fluorogenic protease substrate, showed that cells expressing the canine TMPRSS1 and TMPRSS2 proteases, or the human TMPRSS2 protease, cleaved only a small amount of the substrate, whereas significant cleavage of the substrate was observed by the canine TMPRSS4 protease. Collectively, these findings suggest that, of the proteins investigated, only the canine TMPRSS4 protease is transported to the plasma membrane to a significant extent.

TMPRSS4 transcripts were also detected in the MDCK-MR subline, although these cells did not support the trypsin-independent replication of H1N1_PR8_. One possible explanation for this apparent contradiction is that the TMPRSS4 levels produced in these cells are too low to efficiently activate the virus, or that the protein is expressed in a mutated, non-functional form. Moreover, transcripts for TMPRSS1 and matriptase were detected in MDCK-MR cells. Sequencing and expression studies are required to clarify whether these proteases are expressed in a proteolytically active form. It also remains to be shown whether MDCK-MR cells express serine protease inhibitors such as plasminogen activator inhibitor 1 (PAI-1) (37) or Kunitz-type inhibitors such as hepatocyte growth factor inhibitor 2 (HAI-2) (38) at levels that would suppress endogenously expressed transmembrane serine proteases. It is noteworthy that among the four MDCK sublines tested, the MDCK-MR line was the only one where both TMPRSS2 and HAT transcripts were absent.

The generation of HEK-293T cells constitutively expressing canine TMPRSS2 and TMPRSS4, respectively, allowed us to analyze the proteolytic activation of various IAV strains by these proteases (see **Fig. 10**). We observed that most viral strains were generally activated more efficiently by canine TMPRSS4 than by canine TMPRSS2. The only exceptions were the two H5N1 strains examined, which replicated at significantly higher titers on HEK-293T-TMPRSS2 than on HEK-293T-TMPRSS4 cells. One possible explanation for the more efficient activation of IAV by the TMPRSS4 protease could be that this protease is found in greater quantities on the plasma membrane than the TMPRSS2 protease. There is evidence that TMPRSS2 and other transmembrane serine proteases cleave HA during its transport through the secretory pathway (39, 40). However, it is conceivable that this cleavage in the secretory pathway is only partial and depends on the transport rate of HA. On the other hand, we found that the activation of H1N1_PR8_ by endogenous proteases in MDCK-NBL2 cells was completely inhibited by camostat and nafamostat in the low micromolar range. As both compounds are water-soluble serine protease inhibitors (41), it can be assumed that they acted primarily on proteases that were accessible on the cell surface. This assumption is also supported by earlier reports showing that the cleavage of HA in human respiratory epithelial cells is cell-associated and can be inhibited by serine protease inhibitors such as aprotinin and leupeptin (42).

In addition to the differences in cleavage efficacy between the two proteases, TMPRSS2 and TMPRSS4, significant differences were also observed in the activation of different viral strains by the same protease. For example, H3N2_Bak/03_ replicated in HEK-293T-TMPRSS4 cells without trypsin almost as efficiently as in the presence of trypsin, while H3N2_Ud/72_ replicated significantly less efficiently without trypsin, suggesting that not only subtype-specific but also strain-specific differences in the proteolytic activation of IAV may exist. These differences might be attributable to a different conformation of the HA and, consequently, to different accessibility of the cleavage site loop. In this context, it is important to note that differences in the glycosylation pattern can have a significant influence on the HA conformation and the accessibility of the cleavage site (43–45). On the other hand, site-specific glycosylation of HA can also protect the protein from erroneous proteolytic cleavage (46).

In recent years, TMPRSS2 has attracted a great deal of attention, as it has been shown to be involved in the proteolytic activation of the SARS-CoV-2 spike protein (47, 48). This transmembrane serine protease has been proposed to be the most important protease in primary human respiratory cells capable of activating H1N1, H3N2 and H7N9 virus strains (25). Interestingly, experiments with TMPRSS2^-/-^knockout mice showed that H1N1_PR8_ is unable to replicate in these animals, while H3N2 viruses continue to be proteolytically activated (35). Only a TMPRSS2^-/-^-TMPRSS4^-/-^double knockout prevented the replication of H3N2 virus, suggesting that TMPRSS4 plays an important role in the activation of H3N2 virus in mice (35). In this context, it is also worth mentioning that TMPRSS4 is highly expressed across various cell types of the human respiratory tract, and in some even stronger than TMPRSS2 (25, 40).

The tissue-specific expression of transmembrane protein proteases can have an important impact on viral tissue tropism and pathogenicity (49, 50). Although human influenza viruses primarily replicate in the respiratory tract, replication in extrapulmonary tissues has been reported as well (51, 52). Recently, IAV has been shown to replicate productively in primary human tubular kidney cells and induce factors and mechanisms related to regulated cell death and renal pathology observed in virus-infected patients (53). Likewise, SARS-CoV-2 causes acute kidney injury by directly infecting renal tubules (54). In line with these observations, TMPRSS2 expression has been detected in human kidney (55), where it may be involved in the proteolytic activation of the epithelial sodium channel ENaC in the distal nephron (56). MDCK cells are widely used as a model of renal epithelial cell polarity and transepithelial transport and exhibit several properties characteristic of distal nephron epithelia (57, 58). These cells may represent an interesting cell model to further study the impact of viral infection on renal functions.

In this study, we have shown that the serine protease inhibitors camostat mesylate and nafamostat mesylate can effectively inhibit the activation of IAV in MDCK cells. Interestingly, the therapeutic effect was dependent on the viral dose used. The higher the MOI, the lower the inhibitory effect with a 10 µM concentration of either camostat or nafamostat. This result is surprising since these inhibitors do not act on viral but cellular target proteins. A likely explanation for this phenomenon is that HA competes with the inhibitor for binding to the protease and, in the presence of large amounts of HA, the inhibitor has a lower chance of binding to the active site of the protease. If, therefore, the inhibitors are only administered at a late stage of the infection – that is, once the IAV has already undergone several replication cycles and large quantities of HA have been produced – it is possible that the therapeutic effect will be minimal. If this scenario also applies to the treatment of SARS-CoV-2 infection, it could explain why the use of camostat and nafamostat in clinical trials for the treatment of COVID-19 proved largely disappointing, although other factors – such as low concentrations of the inhibitors reaching the respiratory tract – may have also played a role (59).

Although influenza vaccine are still predominantly produced using embryonated chicken eggs, cell culture-based production using established cell lines such as MDCK cells has attracted considerable interest because it reduces reliance on the egg supply, avoids the selection of egg-adaptive mutations that may compromise antigenic match and vaccine effectiveness, and offers greater manufacturing flexibility (60). However, to guarantee high titer production of influenza vaccines in cell lines, exogenous trypsin is added to the cell culture medium (61). To reduce the risk of potential contamination of the cell cultures by viral pathogens coming from animal-derived trypsin preparations (62), alternative strategies are used including the use of recombinant trypsin or other proteases (63), genetically engineered cell lines permanently expressing human airway proteases such as TMPRSS2 or MSPL (64), or the use of alternative cell lines such as PBS-1 cells that allow IAV propagation in the absence of trypsin (65). Our findings suggest that certain MDCK sublines or clones support trypsin-independent replication of IAV, which could be relevant for influenza vaccine production.

In summary, the present study demonstrates that MDCK cells express several transmembrane serine proteases and support the replication of various IAV strains without the need for exogenous trypsin. Of the proteases examined so far, TMPRSS4 appears to play a prominent role as, unlike other proteases, it is efficiently transported to the plasma membrane and capable of activating many IAV strains. MDCK cells are highly heterogeneous, thus selecting the appropriate subline is crucial for their use in trypsin-independent replication, titration, or vaccine production. Furthermore, MDCK cells are a useful model system for developing and optimizing synthetic serine protease inhibitors, particularly for identifying those that remain active under high viral load conditions.

Future studies should focus on further characterizing individual transmembrane serine proteases, including their human orthologues. It will be interesting to see how the various proteases differ in their activation of IAV and other viruses and whether this influences the tropism of these viruses.

## Materials And Methods

### Cells

Madin Darby canine kidney NBL-2 (MDCK-NBL2) cells were purchased from the American Type Culture Collection (ATCC; CCL-34; Manassas, VA, USA) and maintained in Eagle’s minimal essential medium (EMEM; Pan-Biotech) supplemented with 5% foetal bovine serum (FBS). MDCK-I, MDCK-II, and Caco-2 cells were kindly provided by Georg Herrler (University of Veterinary Medicine Hannover, Germany) and maintained in EMEM with 5% and 10% FBS, respectively. Another MDCK subline was obtained from Michael Matrosovich (University of Marburg, Germany) and was designated MDCK-MR to indicate that were from the Institute of Virology in Marburg. A549 cells were obtained from the European Collection of Authenticated Cell Cultures (ECACC, Porton Down, Salisbury, UK) and grown in Ham’s F12 (Sigma-Aldrich) with 10% FBS. Calu-3 cells were obtained from Silke Stertz (University of Zürich, Switzerland) and maintained in EMEM with 5% FBS. Human embryonic kidney 293T (HEK-293T) cells were obtained from ATCC (CRL-3216) and maintained in DMEM with 10% FBS. Baby hamster kidney 21 (BHK-21) cells were purchased from the German Cell Culture Collection (DSMZ, Braunschweig, Germany) and maintained in Glasgow’s minimal essential medium (GMEM; Gibco) with 5% FBS. BHK-G43 cells, a transgenic BHK-21 cell clone expressing the VSV-G protein in a regulated manner (24), were maintained in GMEM with 5% FBS. All cells were grown at 37 °C in a 5% CO_2_ atmosphere.

### Viruses

The human influenza A viruses A/Puerto Rico/8/1934 (H1N1_PR8_), A/Hamburg/4/2009 (H1N1_HH4/09_), and A/Udorn/307/1972 (H3N2_Ud/72_) were kindly provided by Martin Schwemmle, (University of Freiburg, Germany). A/swine/Bakum/IDT1769/2003 (H3N2_Bak/03_) was provided by Ralf Dürrwald (Robert-Koch-Institut, Berlin, Germany). The low-pathogenic avian influenza viruses A/swan/Potsdam/62/1981 (H7N7_Pot/81_), A/turkey/Italy/1258/05 (H5N2_It/05_), A/teal/Föhr/Wv1378-79/03 (H5N2_Fö/03_), A/duck/Potsdam/1402/86 (H5N2_Pot/86_), A/duck/Potsdam/2216-4/84 (H5N6_Pot/84_), A/duck/Potsdam/619/85 (H5N2_Pot/85_), and A/ostrich/Germany/R5-10/06 (H5N3_Ge/06_) were provided by Timm Harder (Friedrich-Löffler-Institut, Insel-Riems Greifswald, Germany). A/turkey/Wisconsin/1/1966 (H9N2_Wi/66_) was made available by William Dundon (Istituto Zooprofilattico Sperimentale delle Venezie, Legnaro, Italy), A/duck/Hokkaido/Vac-1/04 (H5N1_Hok/04_) is a generous gift from Yoshihiro Sakoda (Hokkaido University, Sapporo, Japan).

H5N1_Tex/24_ (HA_mb_) is a genetically modified A/bovine/Texas/24-029328-02/2024 (H5N1) with a monobasic HA cleavage site (66, 67). H5N1_Yam/04_ (HA_mb_) is a genetically modified A/chicken/Yamaguchi/7/2004 (H5N1), in which the polybasic cleavage site has been converted into a monobasic one (68). Using the 8 plasmid-based reverse genetics system for influenza A viruses (69), the reassortant viruses H5N1_Yam/04_ (HA_Iba/05_), H5N1_Yam/04_ (HA_Nl/14_) and H5N1_Yam/04_ (HA_Hu/05_) were generated by replacing the segment 4/HA of A/chicken/Yamaguchi/7/2004 (H5N1) by the corresponding genome segment of either A/chicken/Ibaraki/1/2005 (H5N2) (Genbank accession number: AB261853), A/chicken/Netherlands/14015526/14 (H5N8) (GISAID accession number: EPI547678), or A/wild duck/Hunan/211//05 (H5N1) (GenBank accession numer: EU329186).

Influenza A viruses were propagated on MDCK-II cells in the presence of 1 µg/mL acetylated trypsin (Merck KGaA, Darmstadt, Germany). The supernatant from the infected cell culture was collected 48 h post infection (p.i.), cell debris was removed by centrifugation (1000 × g, 15 min, 4 °C), and the clarified supernatant, supplemented with 10 % FBS, was frozen in 1-mL portions at −70 °C.

### Virus kinetics and titration

Cells of the indicated cell line were seeded in 6-well cell culture plates (TPP Techno Plastic Products AG, Trasadingen, Switzerland) at a density of 250’000 cells/mL (2.5 mL/well). The following day, the cells were inoculated for 60 min with the respective virus at a multiplicity of infection (MOI) of 0.0001. The cells were washed with sterile phosphate-buffered saline containing Ca^2+^ and Mg^2+^ ions (PBS^+/+^). For the “no trypsin” condition, fresh medium with 1% pen-strep was added to the cells. The cells were incubated for 72 h at 37 °C with 2.5 mL/well of FBS-free cell culture medium with or without 1 µg/mL of acetylated trypsin. Cell supernatant (0.25 mL) was collected at 24, 48 and 72 h p.i. and stored in the presence of 10% FBS at −70 °C.

Infectious virus titers were determined using the focus formation assay (FFA). To this end, MDCK-II cells (200’000 cells/ml in EMEM, 5% FBS and 1% penicillin-streptomycin) were seeded into 96-well cell culture plates (TPP; 100 µL/well). The virus-containing cell culture supernatants were diluted in 10-fold dilution steps (in duplicates) and subsequently incubated with the MDCK-II cells (40 µL/well) for 90 min at 37 °C. Thereafter, 160 µL of EMEM containing 1% methylcellulose and 1% penicillin–streptomycin was added to each well and incubated with the cells for 24 h at 37 °C. The cells were washed with PBS^+/+^, fixed with 4% formalin, and permeabilized with 0.25% (v/v) Triton X-100. Infected cells were visualized by incubating the cells first with a monoclonal mouse anti-NP antibody (1:150; ATCC, HB-65) and subsequently goat anti-mouse IgG antibody conjugated with Alexa Fluor 488 (1:500; Thermo Fisher, Zug, Switzerland; cat. no. A-11001). The fluorescent cells were examined under a fluorescence microscope. The virus titer was calculated and expressed as focus-forming units per milliliter (FFU/mL).

### Serine protease inhibitor studies

The serine protease inhibitors camostat mesylate (Merck KGaA, cat. no. SML0057), nafamostat mesylate (Merck KGaA, cat. no. N0289), and AEBSF (4-(2-aminoethyl)benzenesulfonyl fluoride hydrochloride; Pefabloc; Roche, product no. 11429868001) were used. All inhibitors were dissolved in PBS^+/+^ to obtain a 10 mM solution that was filtrated through 0.4 µm-pore size filters and stored in 100 µL-aliquots at −20 °C. To assess the inhibitory activity, serine protease inhibitors were added at the indicated concentrations to the cell culture medium of H1N1_PR8_-infected MDCK-NBL2 cells (MOI of 0.0001). Aliquots (250 µL) of cell culture supernatant were collected at 24, 48 and 72 h p.i. and infectious virus titers determined by FFA (see above). To determine the impact of virus dose on the antiviral efficacy of serine protease inhibitors, MDCK-NBL2 cells were infected with H1N1_PR8_ at MOIs 1, 0.1, 0.01, 0.001, or 0.0001. Subsequently, the infected cells were incubated with EMEM with or without 10 μM of either camostat or nafamostat mesylate. Cell culture supernatant was harvested at 48 h p.i. and infectious virus titers were determined by FFA.

### Western blot analysis

Confluent monolayers of MDCK-NBL2 cells grown in T75 flasks were inoculated for 1 h at 37 °C with H1N1_HH4/09_ in FBS-free EMEM (MOI of 0.0001). Thereafter, the cells were washed with sterile PBS^+/+^ and incubated with FBS-free EMEM medium optionally supplemented with 1 µg/mL. At 48 h p.i., the cell culture supernatant was collected and cell debris removed by centrifugation (1200 x g, 15 min, 4 °C). Subsequently, virus particles were pelleted from the clarified cell culture supernatant by ultracentrifugation (105’000 x g, 1 h, 4 °C) and solubilized by adding to the pellets 100 µL of preheated (95 °C) sodium dodecyl sulphate (SDS) sample buffer optionally containing 0.1 M of dithiothreitol (DTT). The solubilized proteins were separated by SDS polyacrylamide gel electrophoresis (PAGE) using two 4-12% gradient gels (SurePAGE™; Genscript, Leiden, The Netherlands) and transferred to nitrocellulose membranes by semi-dry blotting. The nitrocellulose membrane was blocked overnight at 4 °C with Odyssey Blocking Reagent (Li-COR Biosciences, Lincoln, NE) and subsequently incubated with a rabbit polyclonal immune serum directed against the HA protein of A/California/04/2009 (H1N1) (Thermo Fisher, cat. no. 16398634). The membrane was washed with PBS containing 0.1% Tween-20 and incubated with IRDye 800CW conjugated goat anti-rabbit IgG (LI-COR Biosciences; cat. no. 926-32211) diluted 1:5000 in PBS. Following several washing steps with PBS/0.1% Tween-20, the blots were scanned with the Odyssey Infrared Imaging system (LI-COR Biosciences, Bad Homburg, Germany).

### Detection of canine transmembrane serine protease transcripts by RT-PCR

Total RNA was extracted from MDCK-II and MDCK-NBL2 cells with the NucleoSpin RNA Plus kit (Macherey-Nagel) and reverse-transcribed into cDNA with the High-Capacity cDNA Reverse Transcription kit (Applied Biosystems; cat. no. 4368814). The cDNA was then amplified by Phusion Hot Start II High-Fidelity PCR (Thermo Scientific; cat. no. F549L) using oligonucleotide primers which were designed based on the target gene sequences deposited at GenBank (**Table 1**). GAPDH, a housekeeping gene, was used as a positive control. The RT-PCR products were analysed by agarose gel electrophoresis with subsequent DNA staining by ethidium bromide.

Reverse transcription of RNA transcripts encoding canine TMPRSS1, TMPRSS2, TMPRSS4, HAT/TMPRSS11D, ST14, and GAPDH and subsequent real-time quantitative PCR (qPCR) were performed on a QuantStudio 5 real-time PCR system (Thermo Fisher Scientific) using the AgPath-ID™ One-Step RT-PCR reagents (ThermoFisher Scientific, cat. no. 4387391) and the primers and probes listed in **Table 3**. The following RT-qPCR protocol was used: reverse transcription at 45 °C for 10 min, denaturation at 95 °C for 10 min, 45 amplification cycles with each cycle consisting of denaturation at 95 °C for 15 sec, primer annealing at 56 °C for 30 sec, and strand elongation at 72 °C for 30 sec. Data were acquired and analyzed using the Design and Analysis Software v1.5.2 (Thermo Fisher Scientific). Expression levels of the canine transmembrane serine proteases were calculated relative to the expression level of GAPDH as follows: 1/ΔCt x 100 = % expression level, where ΔCt is the difference between the cycle threshold value for the respective transmembrane serine protease and the cycle threshold value of canine GAPDH.

**Table 3.**
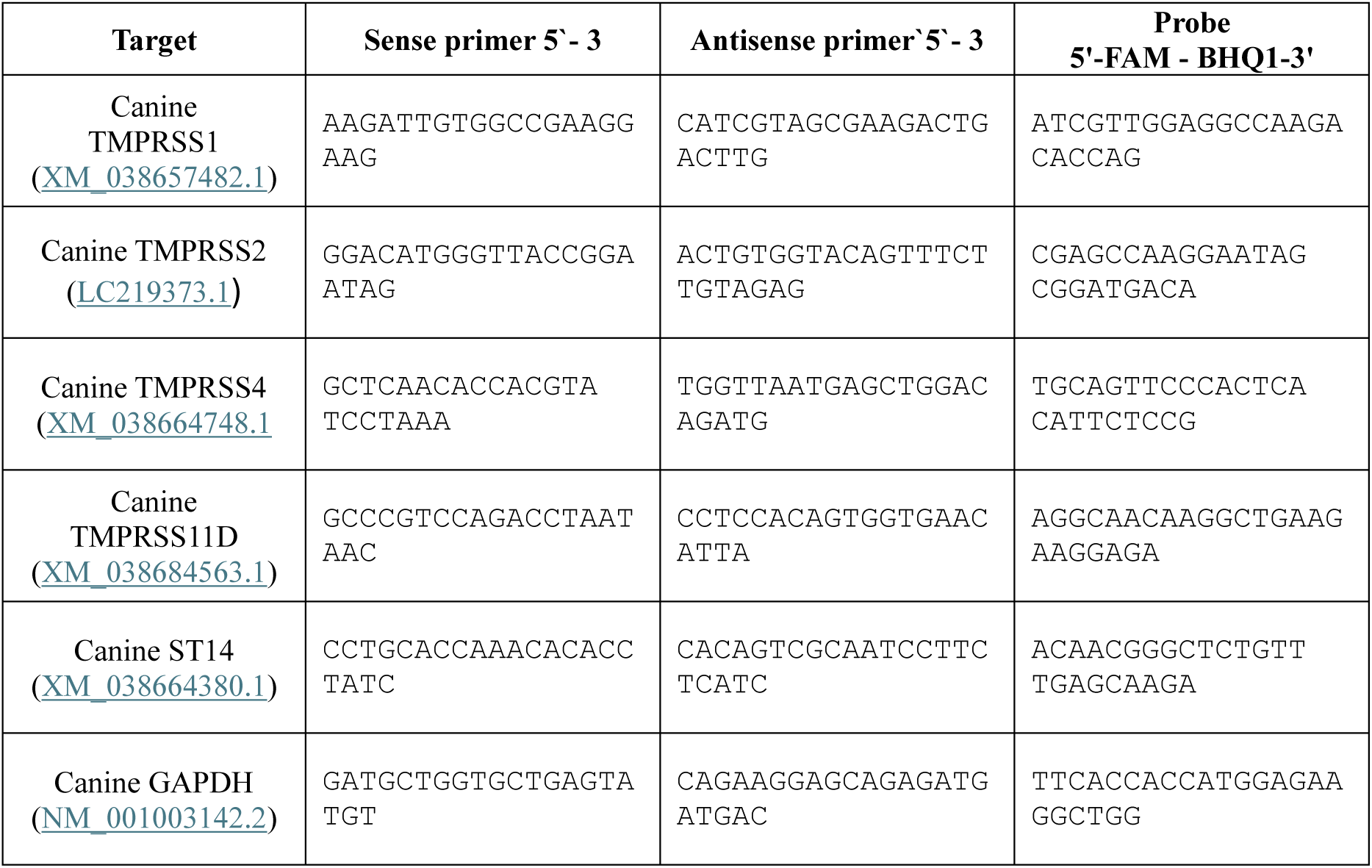
Oligonucleotides for RT-qPCR detection of transmembrane serine proteases.

### Generation of recombinant expression plasmids

Total RNA was extracted from MDCK-NBL2 cells with the NucleoSpin RNA Plus kit (Macherey-Nagel, Düren, Germany, cat. no. 740984.50) and reverse-transcribed into cDNA with the High-Capacity cDNA Reverse Transcription kit (Thermo Fisher Scientific, cat. no. 4368814). The cDNAs of the canine TMPRSS1, TMPRSS2 and TMPRSS4 proteases were amplified by PCR using the Phusion Hot Start II High-Fidelity DNA polymerase (Thermo Scientific) and the primer pairs indicated in **Table 4**. The PCR amplicons were treated for 20 h at 37 °C with the restriction endonucleases indicated in Table 4 and ligated into the correspondingly treated pCEP4 (Thermo Fisher Scientific, cat. no. V04450) and pCMV-3Tag3A plasmids (Agilent Technologies, cat no. 240197), respectively. Human TMPRSS2 cDNA (GenBank accession number NM_001135099) was synthesized by Genscript (Rijswijk, The Netherlands) and cloned into the pCEP4 vectors using the *Nhe*I and *Xho*I endonuclease restriction sites. The complete sequences of the inserts were verified by Sanger sequencing using the BigDye Terminator V3.1 Cycle Sequencing kit (Thermo Fisher Scientific, cat. no. 4337455) and analysed with the SeqStudio Genetic Analyzer and Sequence Scanner software v1.0 (Thermo Fisher Scientific/Applied Biosystems). The consensus sequences of the recombinant genes were obtained by data analysis with Clone Manager 9 software (Sci Ed Software).

**Table 4.**
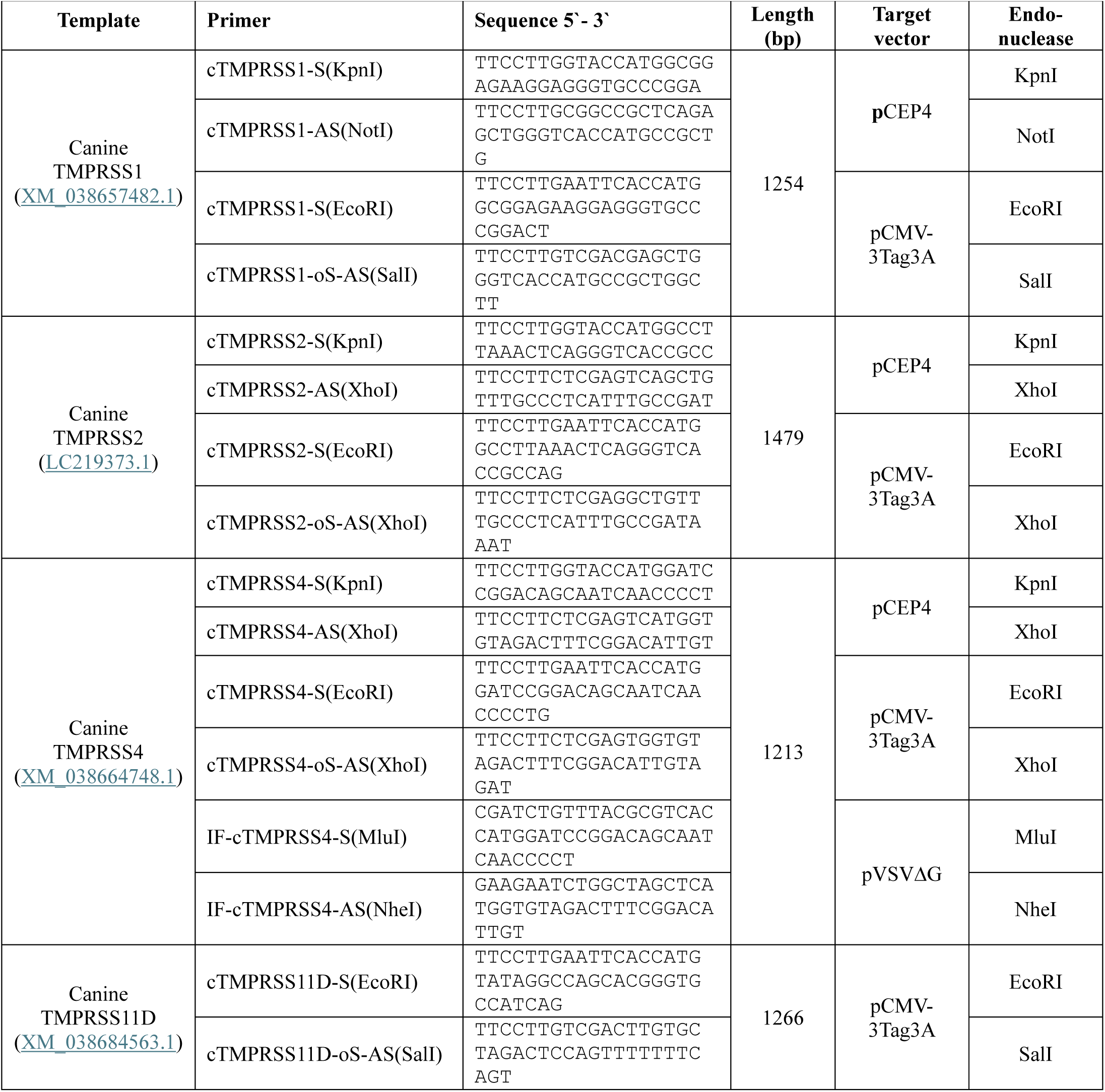
Oligonucleotides for molecular cloning of cDNAs encoding canine transmembrane serine proteases.

### Generation of immune sera using VSV-vectored canine transmembrane serine proteases

The animal experiments were performed according to Swiss laws (the Animal Welfare Act TSchG SR 455, the Animal Welfare Ordinance TSchV SR 455.1, and the Animal Experimentation Ordinance TVV SR 455.163). All experiments were reviewed by the committee on animal experiments of the canton of Bern and were approved by the cantonal veterinary authority under license number BE10/2023.

The cDNAs of canine TMPRSS4 was amplified with the Phusion high fidelity DNA polymerase using the oligonucleotide primers IF-cTMPRSS4-S(*Mlu*I) and IF-cTMPRSS4-AS(*Nhe*I) (**Table 4**) and inserted into the *Mlu*I and *Nhe*I sites of the pVSV*ΔG(HA) plasmid (70) by In-Fusion cloning (In-Fusion Snap Assembly Master Mix, Takara, Saint-Germain-en-Laye, France, cat. no. 638948), resulting in the plasmid pVSVΔG(cTMPRSS4). The whole open reading frame of the cloned canine TMPRSS4 cDNA was determined by Sanger sequencing.

To generate recombinant VSVΔG(cTMPRSS4) virus, BHK-G43 cells were first infected with Modified Virus Ankara encoding T7 RNA polymerase (MVA-T7) and subsequently transfected with the genomic pVSVΔG(TMPRSS4) plasmid along with three plasmids encoding the VSV proteins N, P, and L, respectively (71). The recombinant VSVΔG(TMPRSS4) was rescued from the transfected cell culture supernatant, propagated on BHK-G43 cells, and stored in aliquots at −70 °C in the presence of 10% FBS. Infectious virus titers were determined by end point titration. Infected cells were detected by indirect immunofluorescence using the monoclonal antibody mAb 23H12 directed to the VSV matrix protein (KeraFast, Boston, MA, cat. no. EB0011) and anti-mouse IgG conjugated with Alexa Fluor-488 (diluted 1:500 in PBS; Life Technologies, cat. no. A28175) (72).

For large scale vector production, BHK-G43 cells were seeded into T150 flasks and maintained in 40 mL/flask of GMEM medium with 5% FBS. When confluency was reached, the medium was aspirated and the cells maintained at 37 °C for 6 h with 40 mL/flask of GMEM containing 10^-9^ M of mifepristone. VSVΔG(TMPRSS4) was added (MOI of 0.05), and incubated with the cells for 20 h at 37 °C. The cell culture supernatant was transferred to 50-mL Falcon tubes and cell debris removed by centrifugation (1000 x g, 15 min, 4 °C). The virus particles were pelleted from the cleared supernatant by ultracentrifugation (105,000 x g, 60 min, 4 °C), resuspended in PBS and stored in aliquots at −70 °C prior to vaccination of chickens.

Two specific pathogen-free (SPF) chickens (White leghorn breed; > 1 year of age) were immunized by injection of 0.5 mL of VSVΔG(TMPRSS4) suspension containing 2x 10^8^ infectious units/mL into the breast muscle. Four weeks after the first immunization, the animals were boosted with the same vaccine, dose, and inoculation route. Three weeks after the second immunization, the animals were euthanized and bled. The serum was prepared by centrifugation of the coagulated blood and stored in 1-mL aliquots at −20 °C.

### Generation of HEK-293T cells expressing canine transmembrane serine proteases

HEK-293T cells (400’000 cells/mL) were seeded into 10-cm diameter cell culture dishes (10 mL/dish) and transfected with 10 µg of either pCEP4, pCEP4-TMPRSS2, or pCEP4-TMPRSS4 using 20 µL of Lipofectamin 2000 transfection reagent. For the constitutive expression of the canine transmembrane serine proteases, the transfected cells were selected with 100 µg/mL of hygromycin B (Invivogen, Toulouse, France; cat. no. ant-hg-1). In addition, 10 µM of nafamostat mesylate were added to the cell culture medium to avoid cytotoxic effects due to TMPRSS overexpression. Because of the episomal replication of the pCEP4 plasmid in human cells, the whole hygromycin B-resistant cell population was used for further analysis without the selection of individual cell clones.

### Indirect immunofluorescence analysis

MDCK-MR cells were seeded onto sterile coverslips (12-mm in diameter) that were placed in a 24-well cell culture plate (50’000 cells/well) and incubated for 20 h at 37 °C in MEM medium with 5% FBS. Thereafter, the cell culture medium was aspirated and replaced with 0.6 mL/well of fresh MEM/5% FBS. The cells in each well were transfected by adding 100 µL of OPTI-MEM (Life Technologies, Zug, Switzerland, cat. No. 31985-062) containing 1 µg/well of either pCMV-3Tag3A plasmid, pCMV-cTMPRSS1-3FLAG, pCMV-cTMPRSS2-3FLAG, pCMV-cTMPRSS4-3FLAG, or pCMV-cTMPRSS11D-3FLAG and 2 µL of Lipofectamin 2000 transfection reagent. For each plasmid construct, cells in four wells were incubated with the transfection mix for 20 h at 37 °C, washed once with PBS^+/+^, and fixed for 30 min at 21 °C with PBS containing 4% formalin. The cells were washed twice with PBS^+/+^ containing 0.1 M glycine and once with PBS^+/+^. Cells in two out of the four wells transfected with the same plasmid construct were additionally permeabilized for 5 min at 21 °C with PBS containing 0.25% Triton X-100. The cells were incubated for 1 h at 21 °C with a mouse monoclonal anti-FLAG antibody (clone M2; Merck KGaA, cat. no. F3165), diluted 1:200 in PBS, washed three times with PBS, and then incubated for 1 h at 21 °C in the dark with a goat anti-mouse IgG conjugated to Alexa 488 (Life Technologies, cat. no. A11001). The cells were washed as above and stained for 10 minutes at 37 °C with 4′,6-diamidino-2-phenylindole (DAPI, 0.1 µg/mL in 70% ethanol; Merck KGaA, cat. no. 9542), embedded in Mowiol 4–88, and analyzed by fluorescence microscopy (Observer.Z1 microscope, Zeiss, Feldbach, Switzerland).

For immunofluorescence staining of HEK-293T cells stably transfected with either pCEP4 or pCEP4-cTMPRSS4, the cells were first seeded onto sterile coverslips (12-mm in diameter, placed in a 24-well cell culture plate) at 200’000 cells/well, and cultured for 20 h at 37 °C. The cells were incubated for 1 h at 21 °C with PBS^+/+^ containing 4% formalin, washed twice with PBS^+/+^ containing 0.1 M glycine, and once with normal PBS^+/+^. The cells were first incubated for 1 h at 21 °C with chicken anti-cTMPRSS4 immune serum (diluted 1:500 in PBS), washed three times with PBS, and subsequently incubated for 1 h at 21 °C with goat anti-chicken IgY conjugated with Alexa Fluor-488 (1:1500, Invitrogen, A-11039). Finally, the cells were three times with PBS, stained with DAPI, and analysed by fluorescence microscopy as described above.

### Serine protease assay

To determine whether the cloned canine transmembrane serine proteases are proteolytically active, we transfected BHK-21 cells with expression plasmids and incubated them with a synthetic fluorogenic peptide substrate. BHK-21 were used because they can be transfected with high efficacy and showed only very low endogenous serine protease activity. The assay was performed in a 96-well format by seeding 100 µL of a BHK-21 cell suspension (200’000 cells/mL) into each well and incubated them at 37 °C in GMEM medium supplemented with 5% FBS, usually reaching 80 – 90% confluency after 20 h of incubation. The cells in each well were transfected with 0.1 µg of plasmid DNA encoding the corresponding transmembrane serine protease and 0.2 µL of Lipofectamin 2000 transfection reagent (Life Technologies, Zug, Switzerland, cat. no. 11668019). The cells were incubated for 20 h at 37 °C, washed once with FBS-free GMEM medium and further incubated in this medium for 4 h. The transfected cells were subsequently incubated 2 h, 37 °C with 70 µL/well of PBS containing 0.9 mM Ca^2+^, 0.49 mM Mg^2+^ and 50 µM of the serine protease substrate BOC-QAR-AMC (R&D Systems, Minneapolis, MN, USA; cat. no. ES014). After the incubation, 60 µL of the inoculum was transferred to black 96-well microtiter plates and fluorescence recorded at 365 nm for excitation and 414/445 nm for emission using a GloMax microplate reader (Promega, Dübendorf, Switzerland).

## Acknowledgements

We are grateful to Timm Harder (FLI, Riems-Greifswald, Germany), Yoshihiro Sakoda (Hokkaido University, Sapporo, Japan), Martin Schwemmle (University of Freiburg i. Br., Germany), Ralf Dürrwald (RKI, Berlin, Germany), and William Dundon ( Istituto Zooprofilattico Sperimentale delle Venezie, Legnaro, Italy) for providing influenza A viruses, and to Georg Herrler (University of Veterinary Medicine, Hannover, Germany), Wolfgang Garten (University of Marburg, Germany), and Silke Stertz (University of Zurich, Switzerland) for providing cell lines. We also would like to thank Alexandre Genoud and Nicolas Ruggli (IVI Mittelhäusern, Switzerland) for their support in the production of immune sera in chickens.

